# Increased *APOEε4* expression is associated with reactive A1 astrocytes and the difference in Alzheimer Disease risk from diverse ancestral backgrounds

**DOI:** 10.1101/2020.03.09.983817

**Authors:** A. J. Griswold, K. Celis, P. Bussies, F. Rajabli, P. Whitehead, K. Hamilton-Nelson, G. W. Beecham, D. M. Dykxhoorn, K. Nuytemans, L. Wang, O. K. Gardner, D. Dorfsman, E.H. Bigio, M. Mesulam, S. Weintraub, C. Geula, M. Gearing, E. Martinez-McGrath, C.L. Dalgard, W. K. Scott, J. L. Haines, M.A. Pericak-Vance, J. I. Young, J. M. Vance

## Abstract

*APOEε4* African local genomic ancestry (LA) confers less risk for Alzheimer disease (AD) relative to European LA (LA) carriers. Single nucleus RNA sequencing from AD-*APOEε4/4* frontal cortex found European LA carriers have a 1.45-fold greater *APOEε4* expression (p< 1.8 E10^−313^) and are associated with a unique A1 reactive astrocyte cluster. This suggests a potential mechanism for the increased risk for AD seen in European LA carriers of *APOEε4*.

## Main

Alzheimer disease (AD) is the most common form of dementia^1^. The *APOEε4* allele is the strongest common genetic risk factor for AD^2^. However, the risk for AD conveyed by the *APOEε4* allele varies between populations, with a stronger risk for *APOEε4* homozygotes in Non-Hispanic Whites (NHW) (Odds Ratio (OR) ~15)^3–6^ than for African-Americans (AA) (OR~8) and Africans (OR~3 for)^3–9^. Recent studies have shown the lower risk in African and AA *APOEε4* carriers is associated with the African local genomic ancestry (LA) around the *APOEε4* allele^10^. As there are no distinct amino acid changes in *APOEε4* between AA and NHW, we hypothesized that non-coding variants affecting gene expression are likely involved.

To identify a protective factor in AA LA that could provide insight into potential therapeutic interventions, we obtained frozen frontal cortex tissue (Brodmann area 9) from four AA and four NHW Alzheimer patients, who were homozygous carriers of the African or European LA *APOEε4/4* genotype (Supplementary Table 1) respectively. Single nucleus RNA-sequencing (snRNA-seq) was performed to dissect tissue complexity. After quality control, we obtained data from a total of 47,113 total nuclei (3,112-8,599 nuclei per sample), sequenced at a median depth of ~131,000 reads per cell with ~1800 genes/nucleus (Supplementary Table 2). Figure 1a shows a two-dimensional UMAP plot showing cellular heterogeneity in 42 distinct clusters for the combined eight samples. The Allen Brain Map (https://celltypes.brain-map.org/rnaseq/human/cortex) (Supplementary File 1) and Single-cell atlas of the Entorhinal Cortex in Human Alzheimer’s Disease^11^ (http://adsn.ddnetbio.com) were used to identify cell types. Expression of defining genes for each cluster is shown in Supplementary Figure 1, cell grouping in a UMAP plot in Figure 1b, and a list of cell types in Supplementary Table 3.

**Figure 1.**
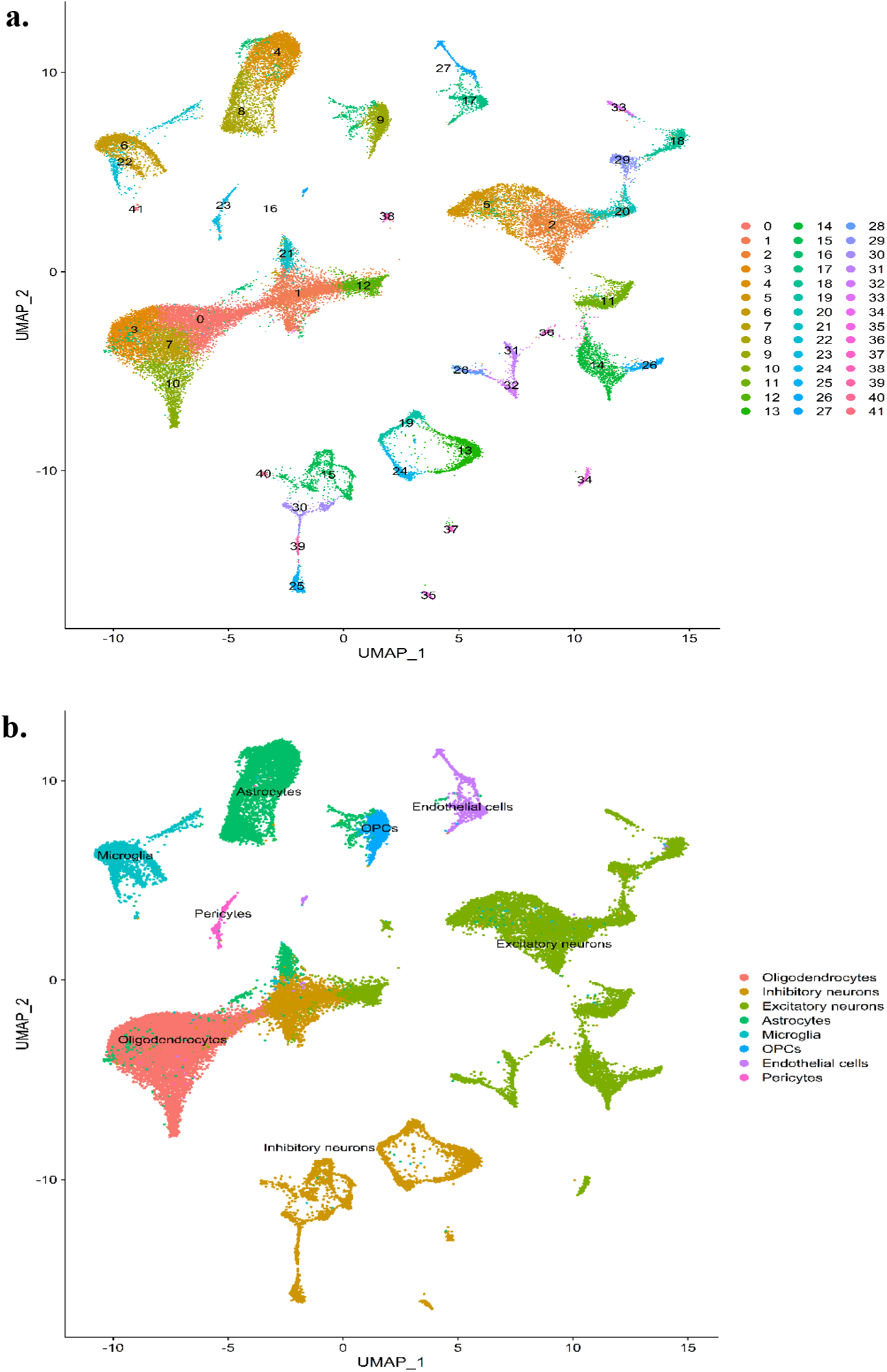
a) UMAP reduction plot of 47,113 nuclei from frontal cortex; *n* = 8 patients. b) UMAP reduction plot showing cell type by color designated in legend and by labels.

*APOEε4* expression was significantly higher overall in EU LA compared to AA LA (adjusted p-value < 1.8E^−313^) and significantly increased in 17 of the 42 clusters (Figure 2 and Supplementary Table 4). Importantly, the expression of *APOEε4* was lower in all four African LA samples, supporting the biological importance between the two groups. Only seven other out of 220 genes within 2Mb on either side of *APOE* had significantly differential expression between AA and NHW in any cluster: *CEACAM19*, *ERCC1*, a long noncoding transcript AC004784.1, *CALM3*, *PLAUR*, *ARHGAP35*, and *HIF3A* (Supplementary Table 5). Of these, none have previously been implicated in AD risk, though *CALM3* encodes for a subunit of calmodulin, and neuronal *APOEε4*^12–15^ has been associated with a dysregulated calcium metabolism. However, none had the consistent and robust differential expression seen in *APOEε4*, strongly supporting this as being the important difference between the African and European *APOEε4* LA.

**Figure 2.**
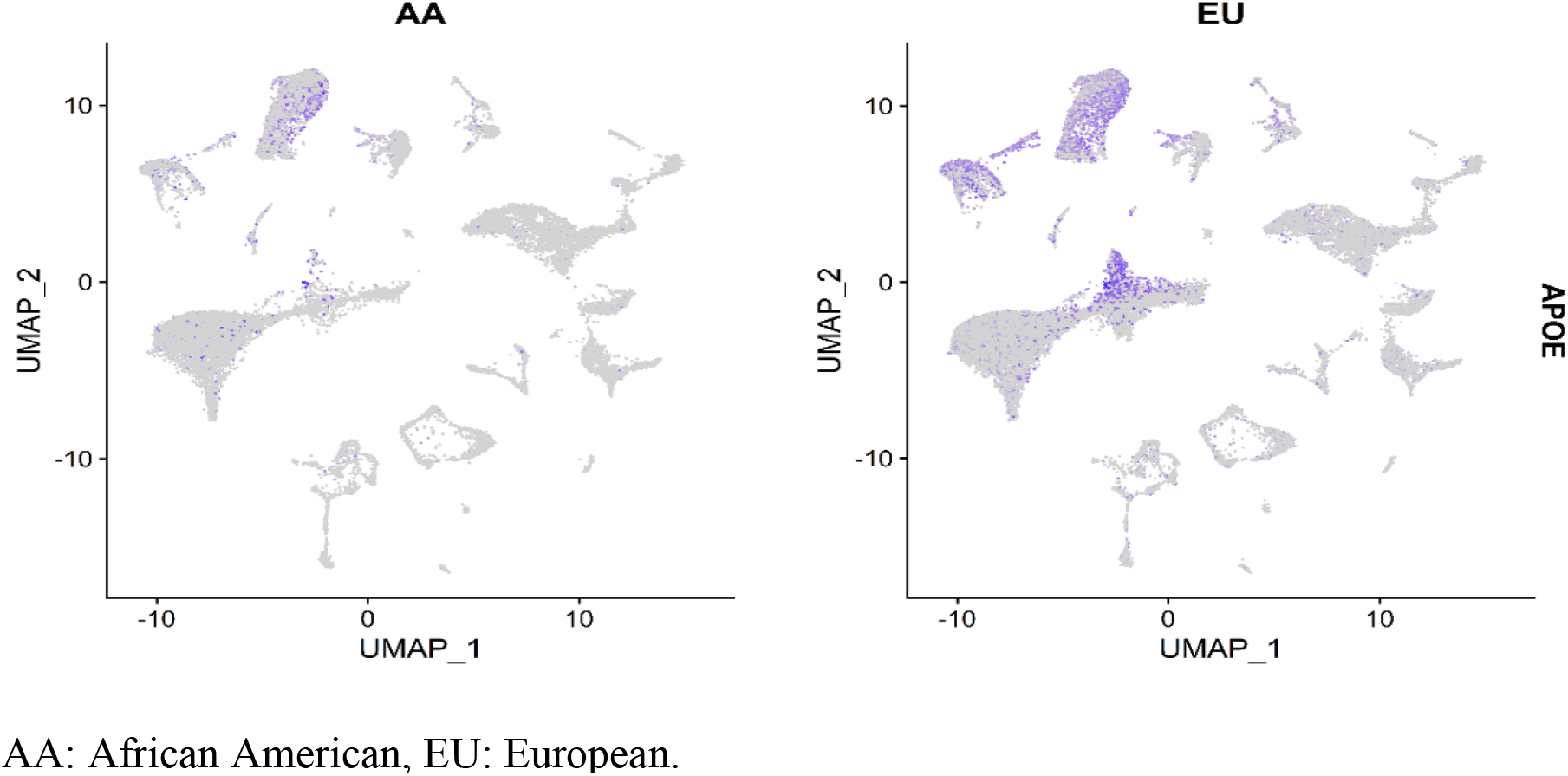
Visualization of *APOE* expression in nuclei from AA local ancestry and EU local ancestry. Cells are overlaid with gene expression information with expression is depicted from gray (low) to purple (high).

We also evaluated differential expression of other known AD genes, including Mendelian (*APP*, *PS1*, *PS2*, and *MAPT*) and genes suggested by GWAS and rare variant association studies in both ethnic groups^16,17^. We observed differential expression between NHW and AA in six reported AD genes including *APP, APOE, CLU, BIN1, PICALM, CR1, PTK2B, SIP1*, and the novel potential AD gene *ADAM10*. Interestingly, we also found significant differential expression in five recently identified novel African-American AD genes^17^: *RBFOX1*, *IGF1R*, *ALCAM*, *GPC6*, and *WDR70* (Supplementary Table 6).

To better understand the molecular pathways altered by ancestry-specific changes in gene expression, we performed an unbiased analysis of enriched pathways. We pooled the DEG between AA and EU LA from all clusters of a given cell-type (Supplementary File 2) to achieve a more complete cell-type transcriptome representation (Supplementary File 3). Pathways enriched in DEG in neurons (both, in inhibitory and excitatory) include neuronal development, synaptic transmission and cell adhesion. DEG in oligodendrocytes are enriched in pathways related to synaptic transmission, brain development, neuronal migration, calcium signaling and response to stimuli. Microglia DEG were enriched in pathways of brain development and signaling. DEG in endothelial cells are enriched in pathways related to responses to stress and angiogenesis.

The most prominent difference between cellular composition between AA and EU LA was in cluster 21 where the proportion of the total cell number in EU LA is ~12 times greater than in the AA LA samples. As the cell proportions of the majority of the clusters are similar (Supplementary Table 3), this finding is unlikely to be secondary to a technical artifact, despite the brain samples being ascertained from different sources. Examination of the transcriptional signature of cluster 21 revealed a strong enrichment for known astrocyte-specific markers (Supplementary Figure 2). To further characterize cluster 21, we compared the transcriptome of this cluster with the other astrocyte clusters (4, 8 and 16). Analysis of the identified DEG from this analysis indicates that cluster 21 preferentially expressed markers of reactive astrocytes^18,19^, with significantly higher levels of *GFAP*, *VIM*, *LGALS1, FGF2*, and *HSBP1* compared to other astrocyte clusters (Supplementary Table 7). Interestingly, compared to the other astrocyte clusters, Cluster 21 also overexpressed *IFITM3*, *CHI3L1*, and *B2M*, which are specifically upregulated by neuroinflammation in A1-type reactive astrocytes in mice^18^. *CHI3L1* (also known as YKL-40) has been proposed as a biomarker for neuroinflammation in early AD^20,21^ and IFITM3 was shown to be upregulated in response to beta-amyloid treatment^22^.

Consistent with an overrepresentation of A1 reactive astrocytes in the EU LA samples as compared to the AA LA samples, enriched pathways using DEG in the astrocytic lineages (clusters 4, 8, 16, and 21) include “regulation of cellular response to heat” (GO:1900034), “response to unfolded protein” (GO:0006986), “protein refolding” (GO:0042026) and “chaperone binding” (GO:0051087) suggestive of differential cellular stress responses between the ancestries. A1 reactive astrocytes are thought to be activated by neuroinflammatory conditions, creating a “toxic” astrocyte, resulting in destructive actions towards neuronal synapses^23^.

Thus, we propose that the higher *APOEε4* expression of *APOEε4* carriers of European LA contributes to a detrimental increase in reactive A1 astrocytes in NHW carriers of the Eu LA *APOEε4*, leading to increased neuronal stress, and increasing the risk for AD. The lower expression of *APOEε4* in AA with the African LA appears to have less effect in driving astrocytes to the A1 reactive phenotype, resulting in the lower risk for AD secondary to *APOEε4* (Figure 2). How this difference in expression is caused by the European LA is currently not known. It could be secondary to sequence differences between the two LA leading to differences in methylation or affecting enhancer activity. Further, little is known about the similarity of open reading frames or topologically associated domains between the two ancestries in the brain, which could affect expression as well. Recently, a SNP in the promoter of *APOE* has been suggested to be related to the risk difference for AD between NHW and Koreans (rs405509)^9^. However, rs405509 is not significantly different in allele frequency between AA and NHW LAs on the *APOEε4* haplotype. Alternatively, the observed increased expression of *APOEε4* could be secondary to an elevation in A1 reactive astrocytes^18^, but this seems unlikely given the increased *APOEε4* expression in multiple cell types. As reactive A1 astrocytes have been shown to be induced through microglia, it seems likely that the final scenario explaining the difference in AD risk in *APOEε4* carriers with NHW vs. AA LA involves a complex interaction of microglia, astrocytes and the observed increase in *APOE* expression^24^.

While this study was designed to evaluate the differences in African and European local ancestry, it also provides one of the first looks at comparing snRNA-seq data between AA and NHW AD patients. The top ten significant DEG between AA and NHW across all clusters are shown in Supplementary Table 8. It includes the *ABCB1* gene involved in the transport of P-glycoprotein and efflux of β-amyloid (Aβ) from the brain, and the *TNR* gene implicated in neurite outgrowth, neural cell adhesion and modulation of sodium channel function. The DEG of the full set of clusters is shown on Supplementary File 2.

One of the challenges in this study was identifying African-American brains for study. The number of African and AA tissue samples currently available for study is limited relative to NHW samples. Further, as an admixed population, many AA samples had mixed LA or even homozygous Eu LA ancestry surrounding *APOEε4* in individuals who self-reported as AA. Our study and that of Rajabli et. al^10^ demonstrate the value of including AA samples and other ancestries in AD studies, including genomic and autopsy research. It also points out that in admixed populations of the Americas, such as AA and Hispanics, the ancestry surrounding the *APOEε4* allele is an important consideration when calculating an individual’s risk for AD.

In summary, this study suggests that the difference in risk between the European and African LAs is secondary to significant differences in the expression of *APOEε4*. The increased expression is strongly associated with an increase in reactive A1 astrocytes in the EU LA. Reactive A1 astrocytes have been shown to cause neuronal stress and eventual cell death, and have been reported in patients with several neurodegenerative diseases including AD^25–27^. Future studies are needed to identify the mechanism producing this expression difference. Identification of the underlying basis for *APOE* overexpression in NHW could lead to therapeutic interventions aimed to reduce the expression of *APOE4* in its carriers, which could have a major effect on the overall risk for AD in the NHW populations.

## Acknowledgements

This study was supported by the National Institute of Aging (R01 AG059018, R01 AG059018, U01 AG052410, U01 AG057659), the Alzheimer Disease Center (ADC) networks (NIA) (AG054074), the BrightFocus Foundation and Alzheimer Association (AG052410). Genomic and data analysis was provided by the Center for Genome Technology (CGT) from the John P. Hussman Institute for Human Genomics (HIHG) from the University of Miami Miller School of Medicine. The work was also funded by P50 AG0256878 from Emory and P30 AG013854 from Northwestern. We thank Dr. Holly Cukier for helpful discussions of the data; Mr. Benjamin Goldstein for data analysis assistance; and the numerous participants, researchers, and staff from many studies who collected and contributed to the data.

## Author Contributions

J.M.V., J.I.Y., A.G., and K.C conceived the project, designed the study, collected the data, performed single nuclei experiments, analyzed data, and wrote the manuscript. F.R. performed local ancestry analysis. P.B., P.W., K.H-N., and D.D. contributed to data collection, data processing, and quality control and cleaning. G.W.B, K.N., L.W., O.K.G., and D.M.D assisted with data analysis and interpretation. E.M-M. and C.L.D performed whole genome sequencing on the samples. E.H.B., M.M., S.W., C.G., M.G., W.K.S., and M.A.P-V. provided brain samples and/or data and provided valuable input to the manuscript. All authors read and contributed to the final manuscript.

## Data availability

All sequencing data generated and processed single nuclei matrix and feature files have been deposited with the Gene Expression Omnibus under accession code GSEXXXXX.

## Code availability

Analysis code is available at https://github.com/agriswold76/AD_ancestry_snRNAseq.

## Methods

### Sample Source

All patients presented clinically with a progressive dementia consistent with AD, had a confirmed diagnosis of Alzheimer Disease upon neuropathological examination, and were *APOEε4* homozygotes. To identify African ancestry patients, the National Alzheimer Coordinating Center (NACC) was screened for patients who self-identified as AA and were *APOEε4* carriers. Autopsy material for the NACC-selected AA samples were obtained from the Department of Neurology Alzheimer Disease Research Centers (ADRC) at Emory University and Northwestern University. NHW autopsy samples were obtained from the John P. Hussman Institute for Human Genomics (HIHG) autopsy programs for AD. All samples were acquired with informed consent for research use and approved by the institutional review board of each center. Frozen frontal cortex tissue (Brodmann area 9) was used for all analyses.

### Selection of Brain Samples

Review of the NACC database initially identified African-American samples with clinical AD confirmed by pathology and an *APOEε4/4* genotype. Material Transfer Agreements (MTA) were successfully completed with two of the largest listed data sets (total 45 samples) containing African American samples, i.e. Northwestern University and Emory University Alzheimer Disease Research Centers (ADRC). Subsequent evaluation revealed that only 14 AA samples were currently available for study. Those 14 samples were assessed for local ancestry in the *APOE* region. Three of the AA samples were homozygous for European LA, and three were heterozygous for European and African LA, excluding them for the study. Two additional samples were eliminated due to the presence of other identified neurologic abnormalities (e.g. global head injury, glioblastoma) that could affect the analysis. This left four samples (three females, one male) that were found to be homozygous for *APOEε4* and consistent for African local ancestry. The African global ancestry of these four samples ranged from 85 to 92%, with the remaining admixture European.

Four NHW samples (three females, one male) were obtained from the HIHG Brain Bank, of which samples collected prior to 2007 were collected at Duke University. Samples were chosen based on tissue availability, sex match, neuropathology (had no other identified neurologic abnormalities that would affect expression), *APOEε4/4* genotype and European local ancestry. Global ancestry was also assessed for all samples, with the NHW samples having >96% European global ancestry.

All donors included in the study were clinically diagnosed with AD using standard cognitive testing (Clinical Dementia Rating (CDR) or Mini-Mental State Exam (MMSE)), and met the neuropathological criteria of the National Institute on Aging-Alzheimer’s Association for the diagnosis of AD. The four AA donors had an age-of-death ranging from 82 to 86 years, versus 70 to 76 years for the four NHW donors. The BRAAK stage from all samples ranged from IV to VI. Description of the samples is shown in Table 1. Whole genome sequencing revealed absence of mutations in any known Mendelian genes for AD (*PSEN1, PSEN2, APP*, and *MAPT*) as well as absence of known rare variants in *ABCA7*, *TREM2* and *SORL1* in all samples.

### Assessment of Genetic Ancestry

All samples were assessed for both Global (GA) and Local Ancestry (LA) using genome-wide genotyping from either the Expanded Multi-Ethnic Genotyping Array, Illumina 1M-duo (v3), or the Global Screening Array (Illumina, San Diego, CA, USA). Global ancestry was estimated by performing Principal Components Analysis (PCA) using the GENESIS R package^28^. The AA and NHW samples were combined with the Human Genome Diversity Project (HGDP) reference panel representing diverse ancestries. To assess the LA, we phased our datasets (SHAPEIT version 2^29^) using 1000 Genomes Phase 3 (1kGP) reference panel and used RFMix to infer LA at loci across the genome^30^. We defined an initial region (chr19:44,000,000–46,000,000Mb) around the *APOE* locus that was broad enough to include potential enhancers, topological associated domains, and other regulatory factors while narrow enough to ensure contiguous LA blocks ^10^. After selecting the *APOE* LA region, we selected individuals homozygous for European (n=4) and African (n=4) local ancestry haplotypes. All individuals used in the study were confirmed homozygous for the *APOEε4* allele by Sanger and whole genome sequencing.

### Known AD mutation screening

Whole Genome Sequencing (WGS) was performed from 1μg of DNA extracted from the brains using the Illumina TruSeq DNA PCR-Free Kit and sequencing to 30X on the Illumina NovaSeq 6000 at either the Center for Genome Technology at the John P. Hussman Institute for Human Genomics or The American Genome Center at Uniformed Services University of the Health Sciences (USUHS). Resulting FASTQ files were aligned to the GRCh38 human reference genome with BWA^31^, followed by Genome Analysis Toolkit (GATK)^32^ base quality score recalibration, duplicate removal, and joint genotype calling across all eight samples with the GATK Haplotype Caller according to GATK Best Practices recommendations^33, 34^. Variant annotation was performed with ANNOVAR^35^ and each sample screened for known pathogenic variants in AD genes.

### Nuclei isolation

Nuclei were isolated from ~100mg of frozen frontal cortex brain tissue from Brodmann area 9 at the HIHG using the Nuclei Isolation Kit: Nuclei EZ Prep (Sigma, #NUC101). All tissues were homogenized in ice-cold EZ Lysis buffer with a glass-on-glass dounce homogenizer, 10 strokes with the tight pestle, followed by 10 strokes of the loose pestle, followed by 5 min incubation. Centrifuged nuclei (500× g, 5 min and 4 °C) were washed in ice-cold EZ Lysis buffer, and Nuclei Suspension Buffer (NSB; consisting of 1X PBS, 1% BSA and 0.2 U/μl RNase inhibitor (NxGEN #97065-224). Isolated nuclei were resuspended in NSB, filtered through a 70 μm and 40 μm cell strainer and pellet re-suspended in 2% BSA in PBS. Then, homogenates (2 mL) were layered onto a 1.8M sucrose cushion and ultra-centrifuged at 24,400 rpm at 4°C for 2 hours using a SW28 swinging bucket rotor (Beckman Coulter Optimal centrifuge #L90K). The nuclear pellet was re-suspended in 2% BSA in 1X PBS. After resuspension, nuclei were re-filtered through a 40 μm cell strainer and an aliquot was trypan-blue stained for visual quality assessment, and counted using the Countess Automated Cell Counter (Thermo Fisher).

### Single nucleus RNA sequencing

Single nucleus sequencing was performed in the Center for Genome Technology at the HIHG. Briefly, nuclei at a concentration of 1200 nuclei/mL were loaded on the 10X Genomics Chromium platform to isolate ~7,000 nuclei per sample and create individually barcoded Gel bead-in-Emulsions (GEMs). GEMs were then subjected to reverse transcription to generate unique molecular identified RNA using the Chromium Single Cell 3’ Reagent Version 3 Kit. Sequencing libraries were evaluated for quality on the Agilent Tape Station (Agilent Technologies, Palo Alto, CA, USA), and quantified using a Qubit 2.0 Fluorometer (Invitrogen, Carlsbad, CA). Pooled libraries were quantified using qPCR prior to loading on the Illumina NovaSeq 6000. We targeted 100,000 reads per cell with sequencing parameters suggested by 10X Genomics: Read1, 28 cycles; Index1, 8 cycles; Read2, 98 cycles.

### RNA-seq alignment

We utilized the 10X Genomics CellRanger v3.0.2 software for primary bioinformatics, first creating de-multiplexed FASTQ files from the raw sequencing output with the {-mkfastq} command. The CellRanger {-count} command was then used to map and quantify sequencing reads. Since nuclear RNA includes an abundance of un-spliced RNA, we aligned to a customized GRCh38 reference genome that includes intronic reads in the gene quantification as suggested by the 10X Genomics manual (https://support.10xgenomics.com/single-cell-gene-expression/software/pipelines/latest/advanced/references#premrna) and previous single nucleus sequencing publications^36^.

### Sequencing quality control and integration

Gene expression matrices created by CellRanger were analyzed further with the Seurat 3.1 pipeline^37^ implemented in R v3.6.1 and RStudio v1.2.1335_64× for data filtering, normalization, integration, and downstream analysis. First, we removed poor quality nuclei or potential doublets by removing nuclei with fewer than 200 or greater than 8000 total genes. Next, nuclei with greater than 10% mitochondrial reads were removed to exclude nuclei with excess associated mitochondria. Global-normalization was then performed on all cells from sample independently using the {NormalizeData} function in Seurat to normalize gene expression of each cell by total expression per sample, followed by log transformation. Seurat’s {FindVariableFeatures} function was used to identify the top 5000 variable genes in each sample.

Data integration was performed to identify shared cell states across different samples following a recently published protocol^38^. First {FindIntegrationAnchors} was used to identify ‘anchors’ between samples that represent pairwise correspondences between individual cells that likely originate from the same cell type. Then, these ‘anchors’ were used to harmonize the samples using the {IntegrateData} command to create a fully integrated R object. Finally, global-scaling of the integrated data set to remove batch effects and unwanted sources of biological variation was performed using {ScaleData}.

### Single nucleus clustering

The scaled integrated object was used for further downstream processing to identify common cell types and enable comparative analyses across the samples. First, the first 50 principal components were calculated on the scaled data and the PCs were projected into two dimensions using the UMAP algorithm^39^. Similar cells were clustered from the principal components with the {FindNeighbors} command and clusters defined using a resolution of 1.2 in the {FindClusters} command resulting in 42 distinct clusters.

Finally, {FindConservedMarkers} was used to identify canonical marker genes for each cluster conserved across samples by performing differential gene expression between clusters and combining the p-values using meta-analysis methods from the MetaDE R package.

### Differential gene expression

Comparisons of cellular composition (e.g. counts and proportions of cells of a certain cluster within each group) was investigated using the {table} command. Furthermore, differentially expressed genes (DEG) between conditions and within each cluster was identified using the {FindMarkers} function using the MAST test which employs a generalized linear model framework using cell detection rate across groups as a co-variate^40^.

### Cluster identification

Each cluster was assigned to a cell class on the basis of expression of marker genes from two sources, 1) the Allen Brain Map (https://celltypes.brain-map.org/rnaseq/human/cortex) (Supplementary File 1) and 2) the website developed by the Polo laboratory^11^(http://adsn.ddnetbio.com). Identification of the reactive A1 astrocyte cluster used available literature characterizing transcriptome of that cell type^18, 19, 41^

### Pathway analysis of transcriptome data

Pathway analysis was performed using the gene enrichment analysis tools DAVID^42^ and Enrichr^43,44^. As input lists we used: (i) genes differentially expressed in African vs European LA samples in each identified cell cluster, (ii) genes differentially expressed in African vs European LA samples in multiple clusters representing similar cell types, and (iii) genes differentially expressed in African vs European LA samples in all clusters. Cell type specific background gene lists were obtained from the Allen Brain Map and converted to ensemble IDs to be used as input for DAVID. Enrichment of genes in KEGG pathways and Gene Ontology Biological Process and Molecular Function libraries was determined. An FDR adjusted p-value of 0.05 was used as a cut-off to determine significant enrichment of a pathway.

## Supplementary Figure Legends

**Supplementary Figure 1.**
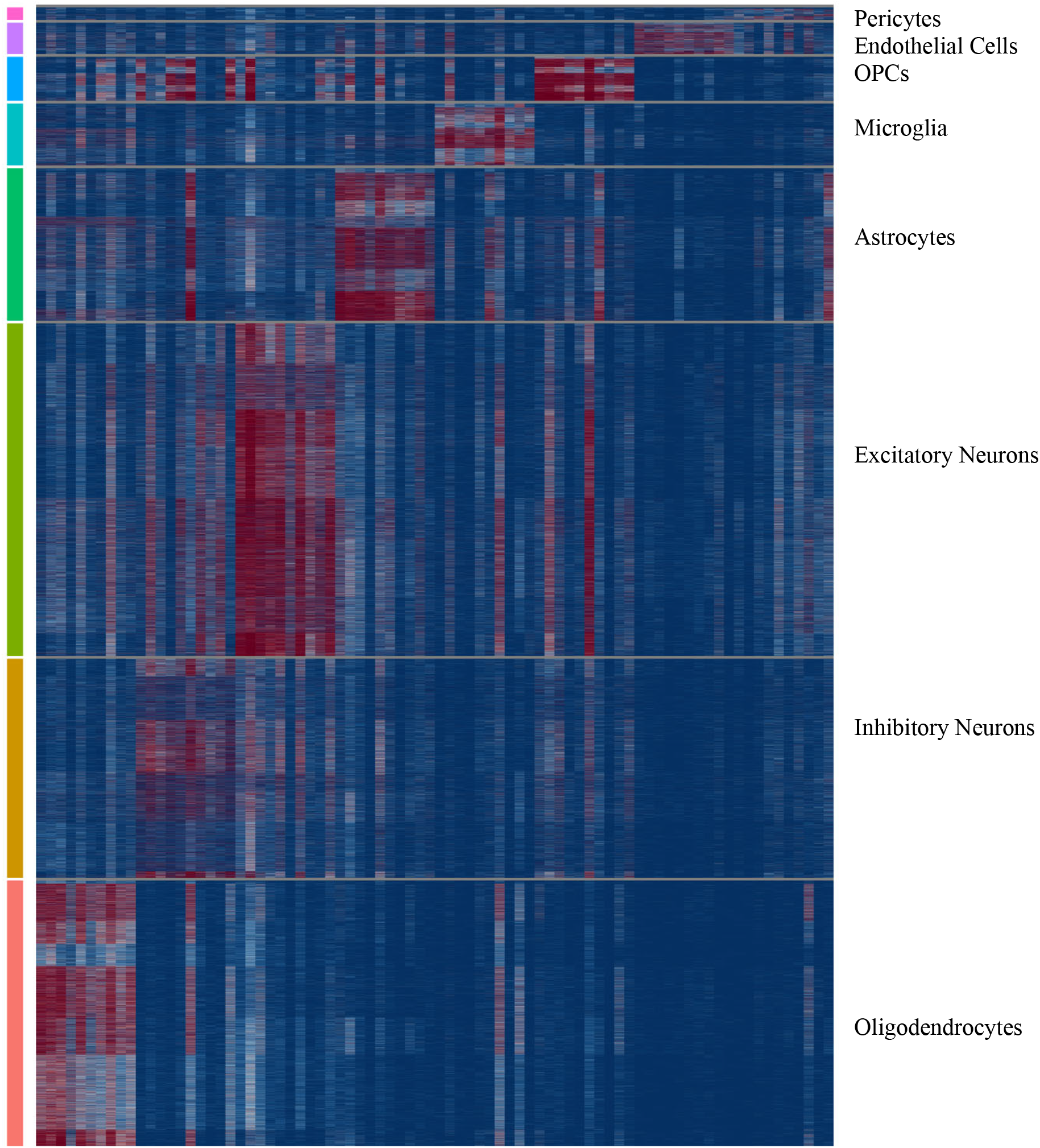
Heatmap of top 10 marker genes defining each cell type.

**Supplementary Figure 2.**
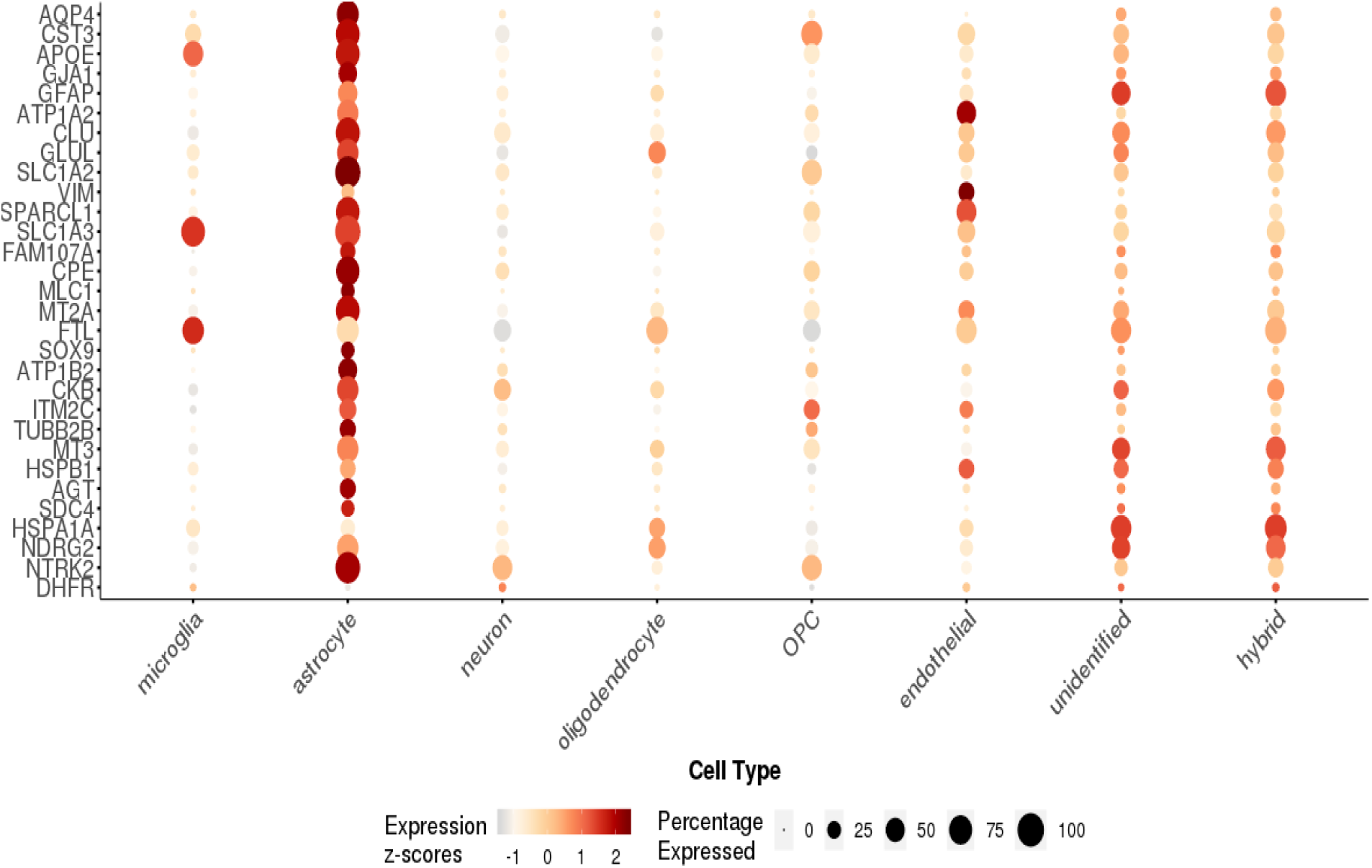
Bubble plot of the top 30 marker genes for cluster 21 (significant adjusted p-value and positive fold change when compared to all other clusters) generated at http://adsn.ddnetbio.com

## Supplementary File Legends

**Supplementary File 1.** Single cell Allen Brain Map (https://celltypes.brain-map.org/rnaseq/human/cortex) heatmap for the top 20 marker genes for each cluster (one per page) used to identify cell types.

**Supplementary File 2.** List of all differentially expressed genes between AA and EU ancestry within each cluster. avg_logFC is the log base 2 fold change. p_val_adj is the MAST FDR corrected p-value.

**Supplementary File 3.** Pathway enrichment analysis in KEGG and GO Ontology Biological Process for the differentially expressed genes from all clusters of a given cell-type. Each tap represents a different cell type.

