## Supplementary File 1 for "Increased *APOEε4* expression is associated with reactive A1 astrocytes and the difference in Alzheimer Disease risk from diverse ancestral backgrounds"

### Supplementary File 3. The Allen Brain Map cell type heatmaps by cluster.

#### Cluster 0

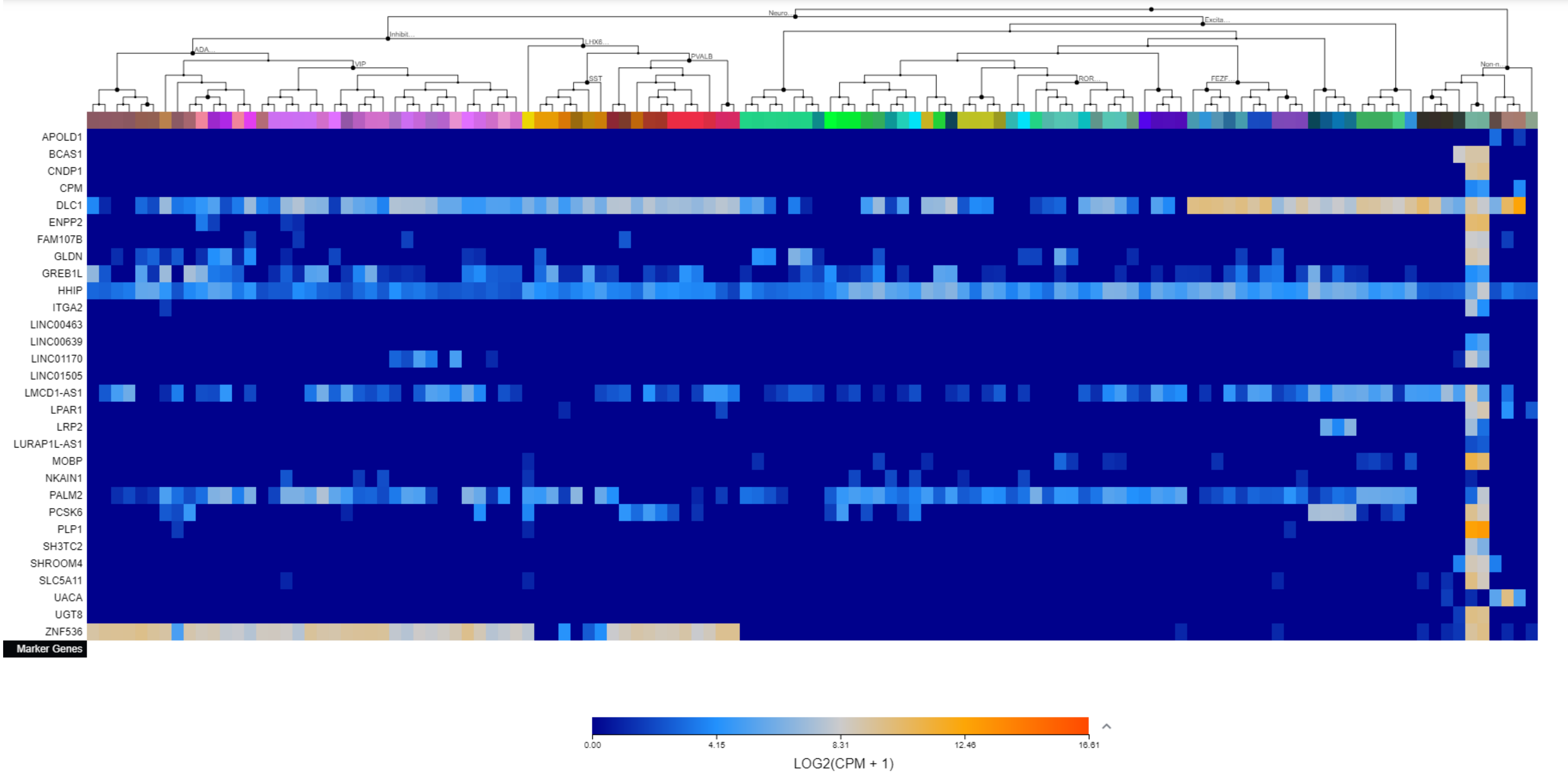

### Supplementary File 3. The Allen Brain Map cell type heatmaps by cluster.

#### Cluster 1

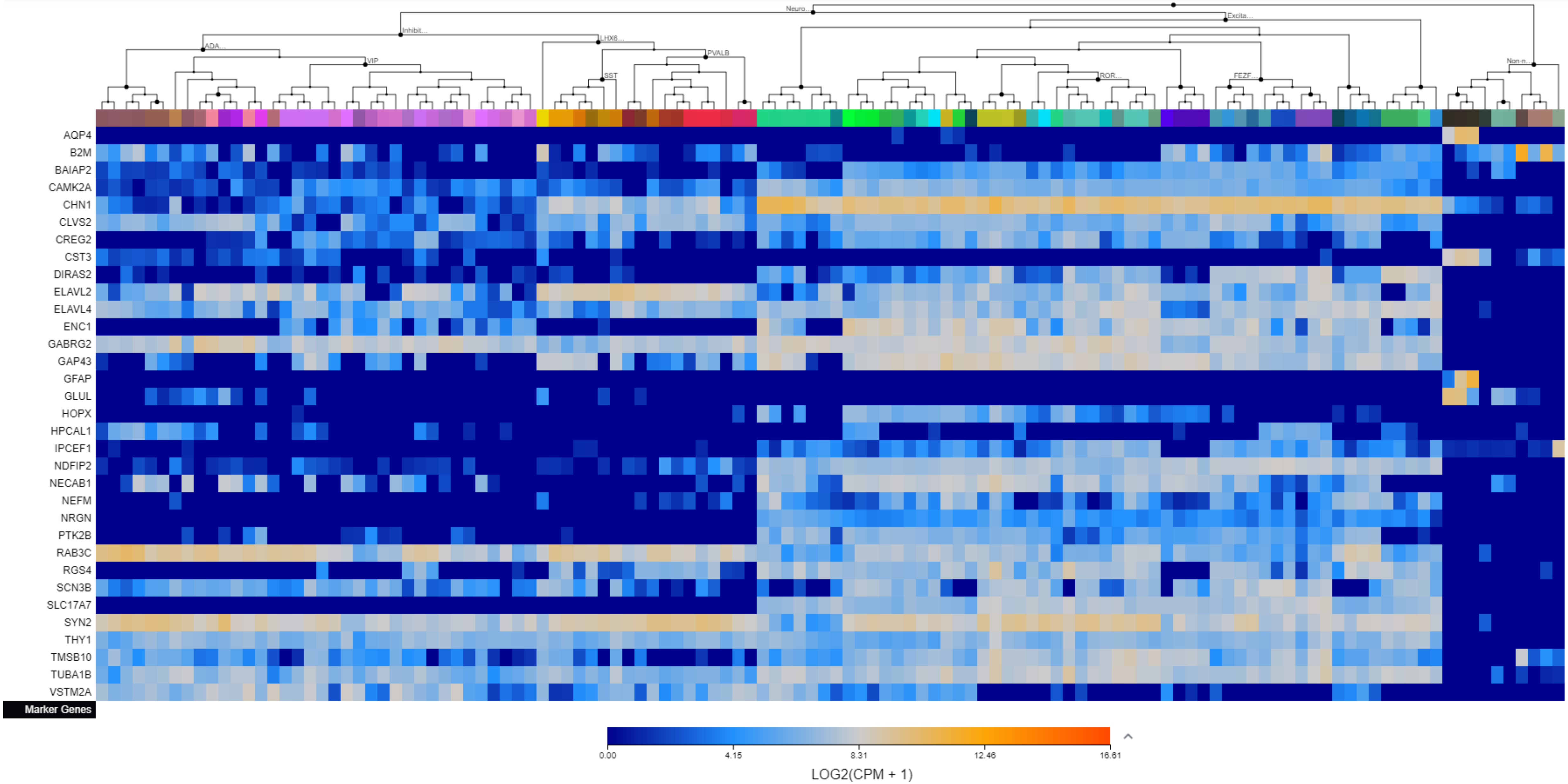

Supplementary File 3. The Allen Brain Map cell type heatmaps by cluster.

Cluster 2

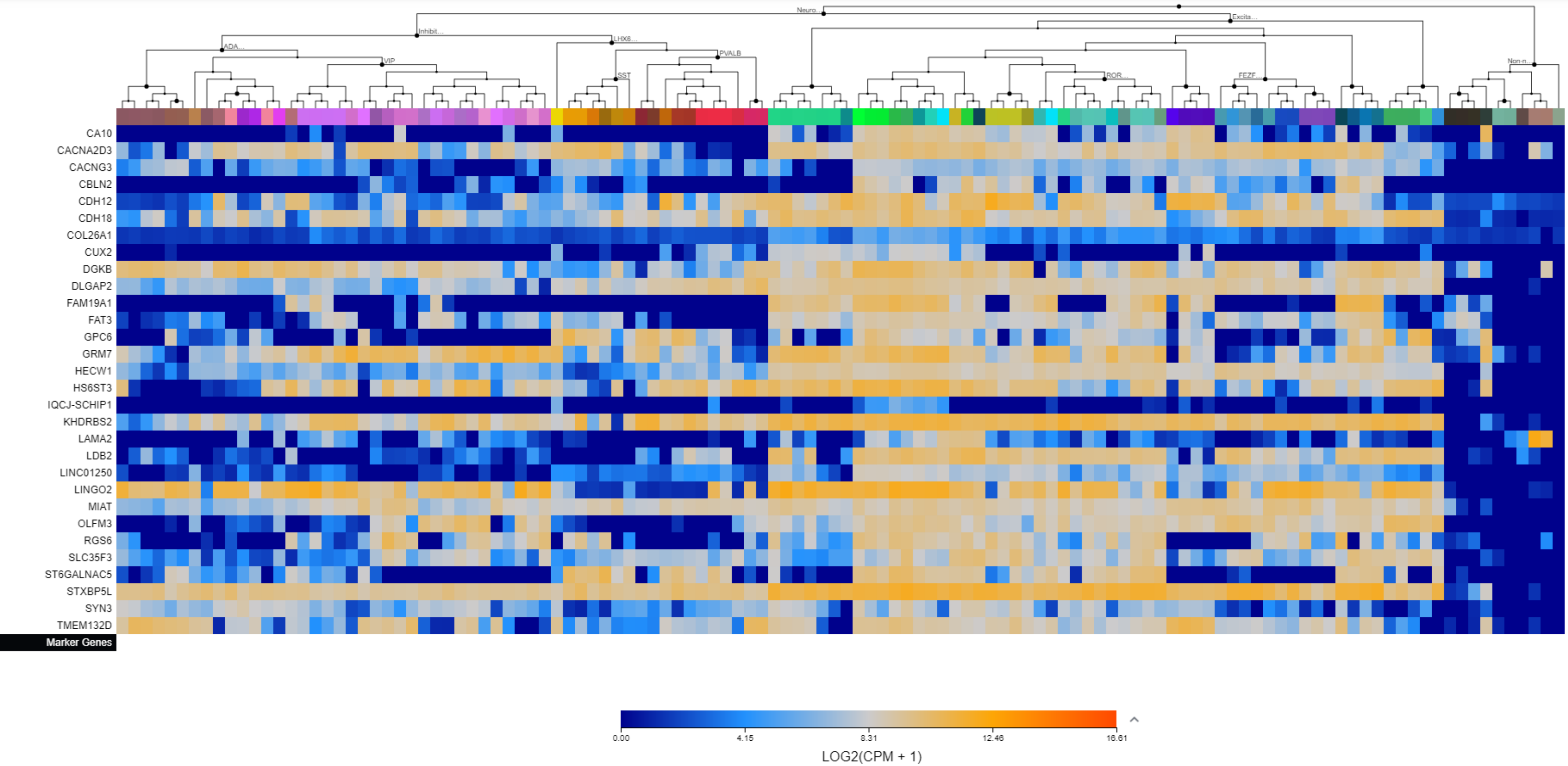

Supplementary File 3. The Allen Brain Map cell type heatmaps by cluster.

Cluster 3

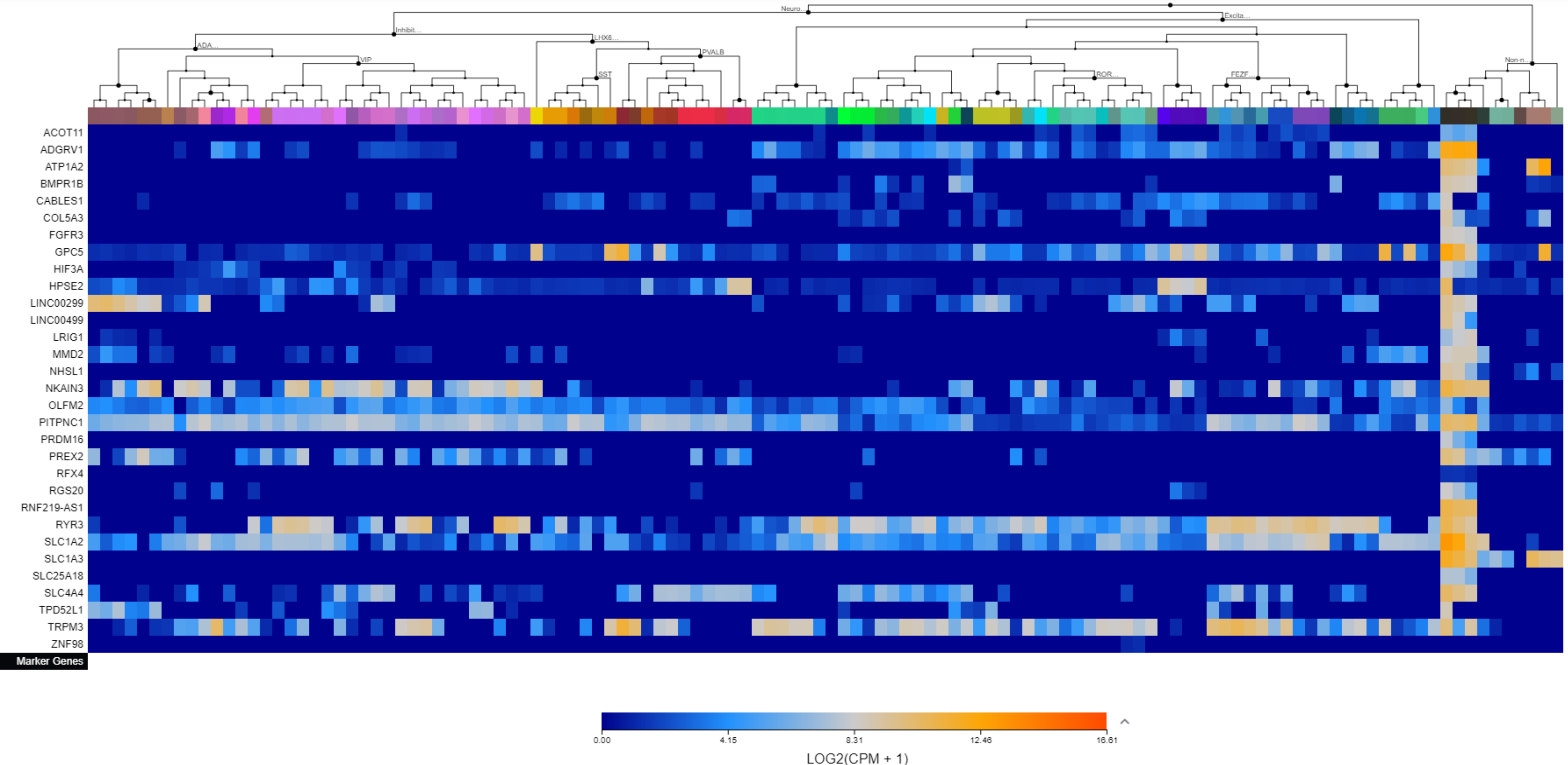

Supplementary File 3. The Allen Brain Map cell type heatmaps by cluster.

Cluster 4

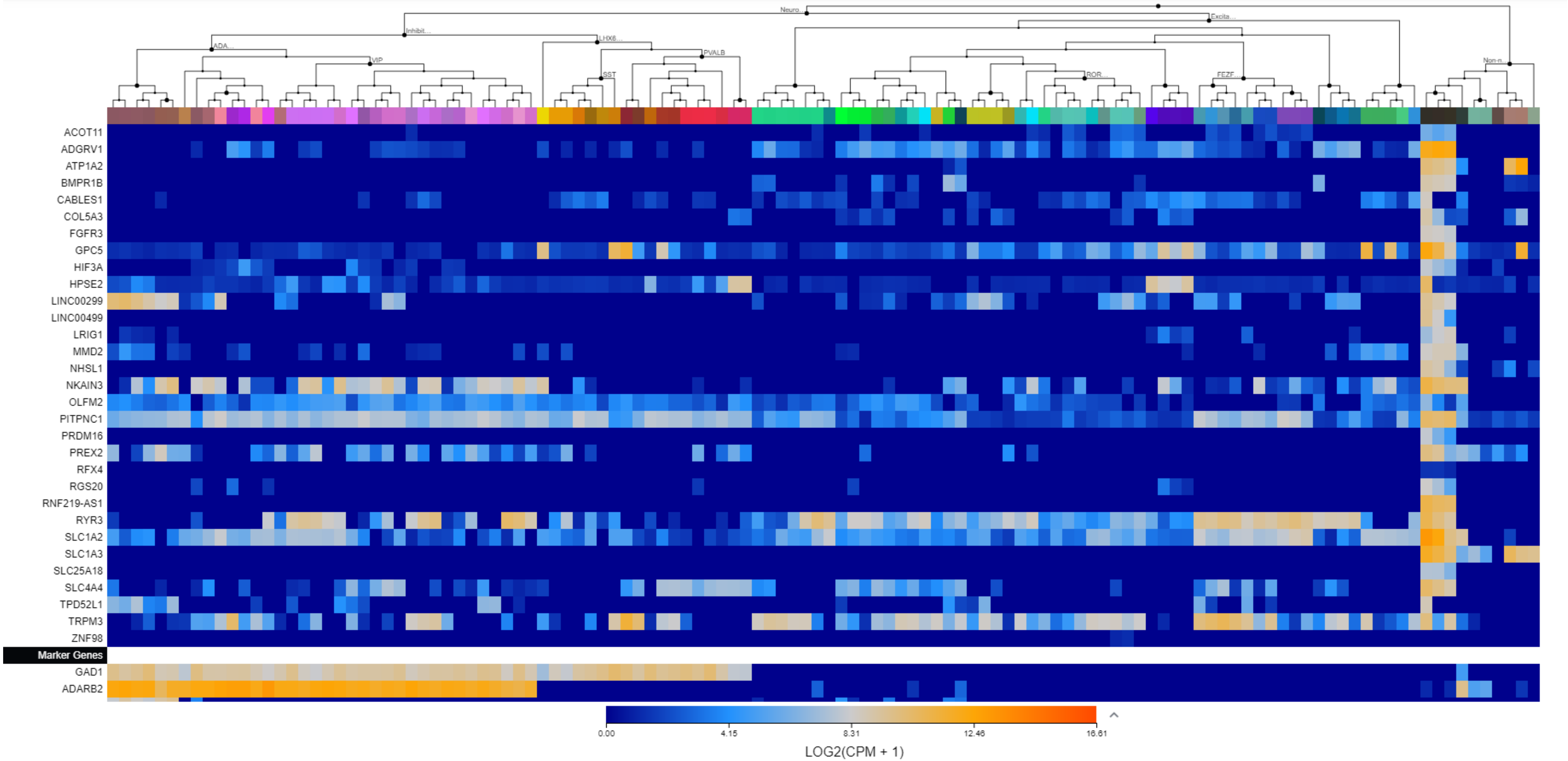

Supplementary File 3. The Allen Brain Map cell type heatmaps by cluster.

Cluster 5

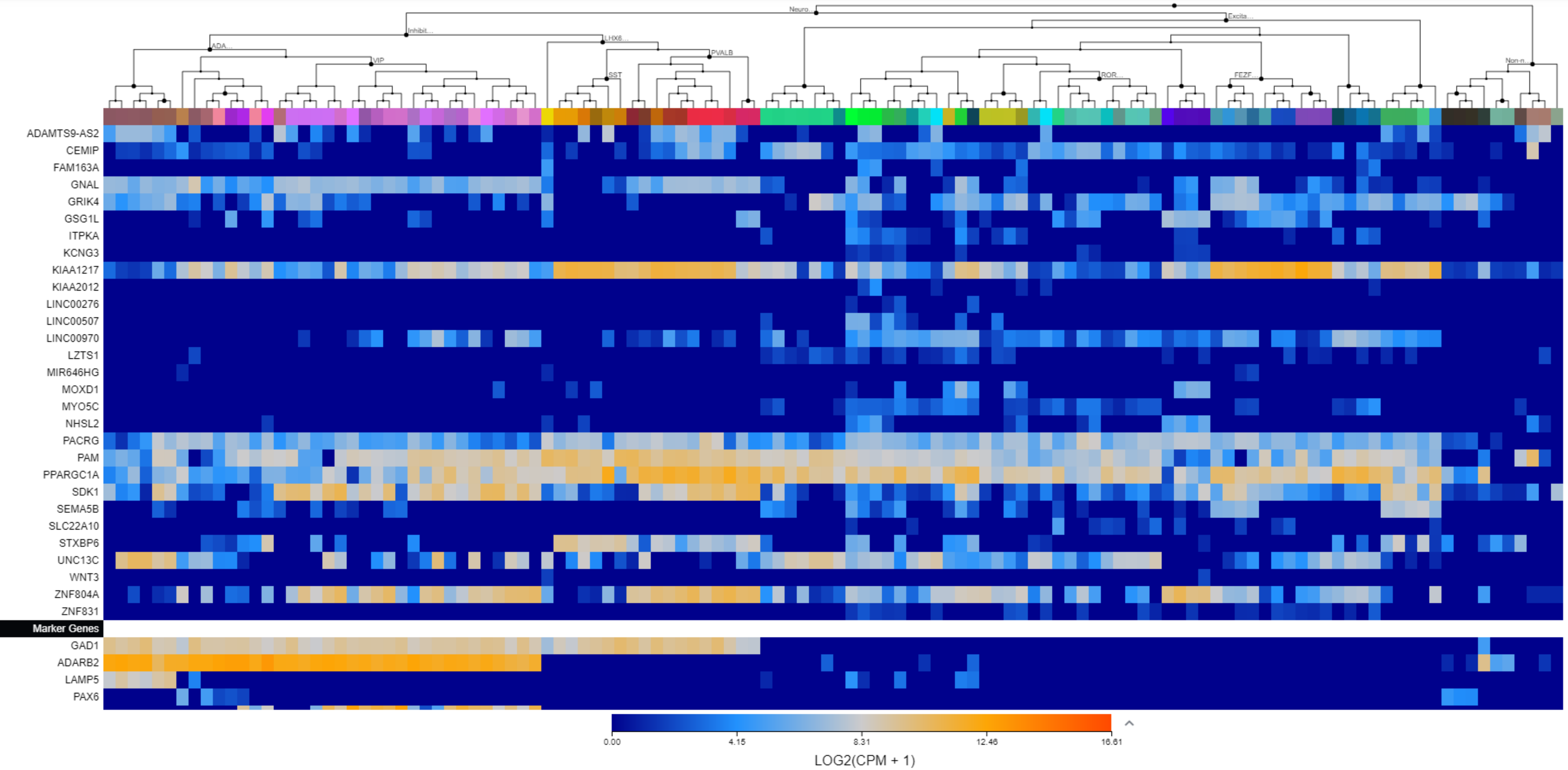

Supplementary File 3. The Allen Brain Map cell type heatmaps by cluster.

Cluster 6

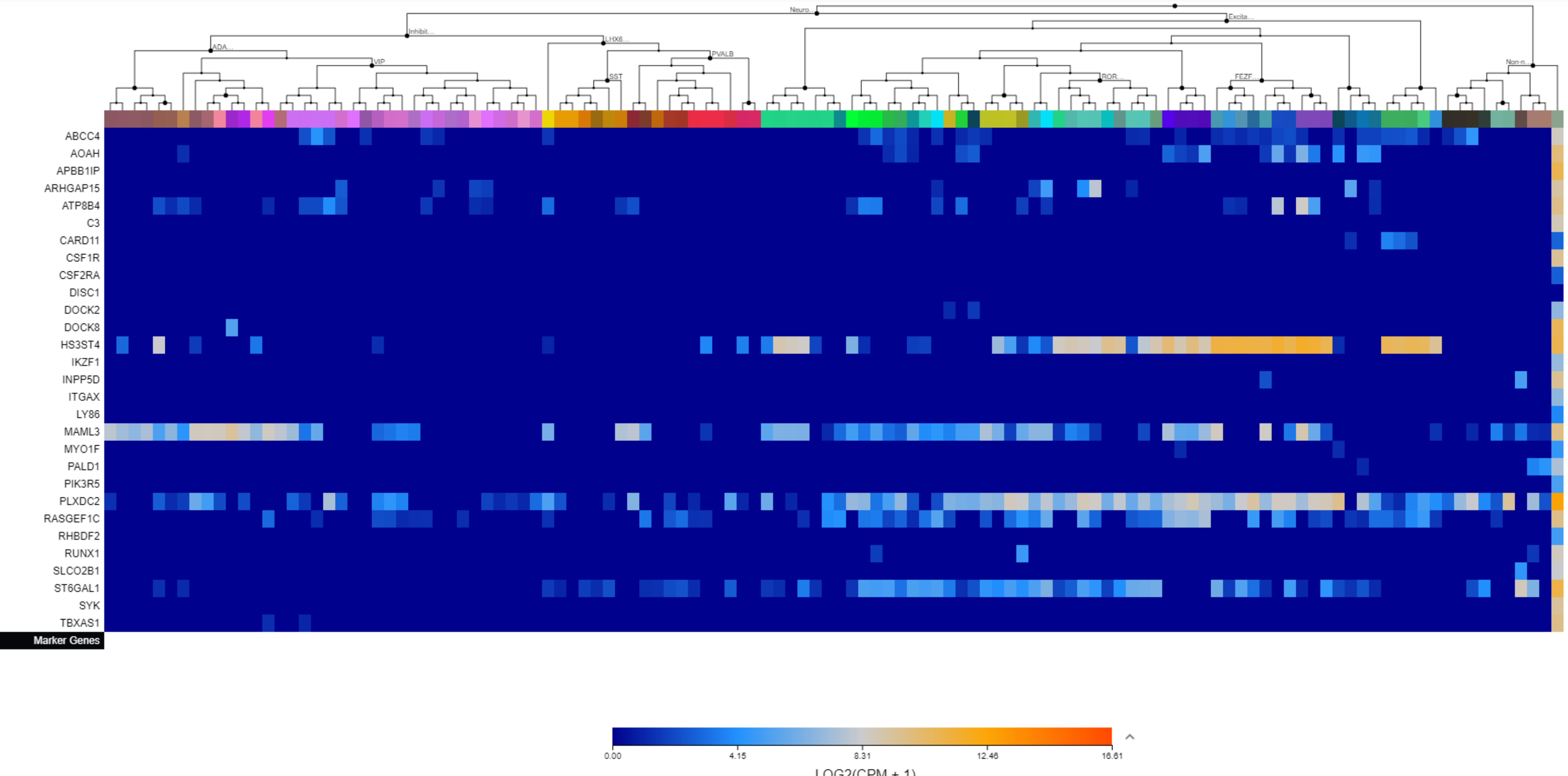

Supplementary File 3. The Allen Brain Map cell type heatmaps by cluster.

Cluster 7

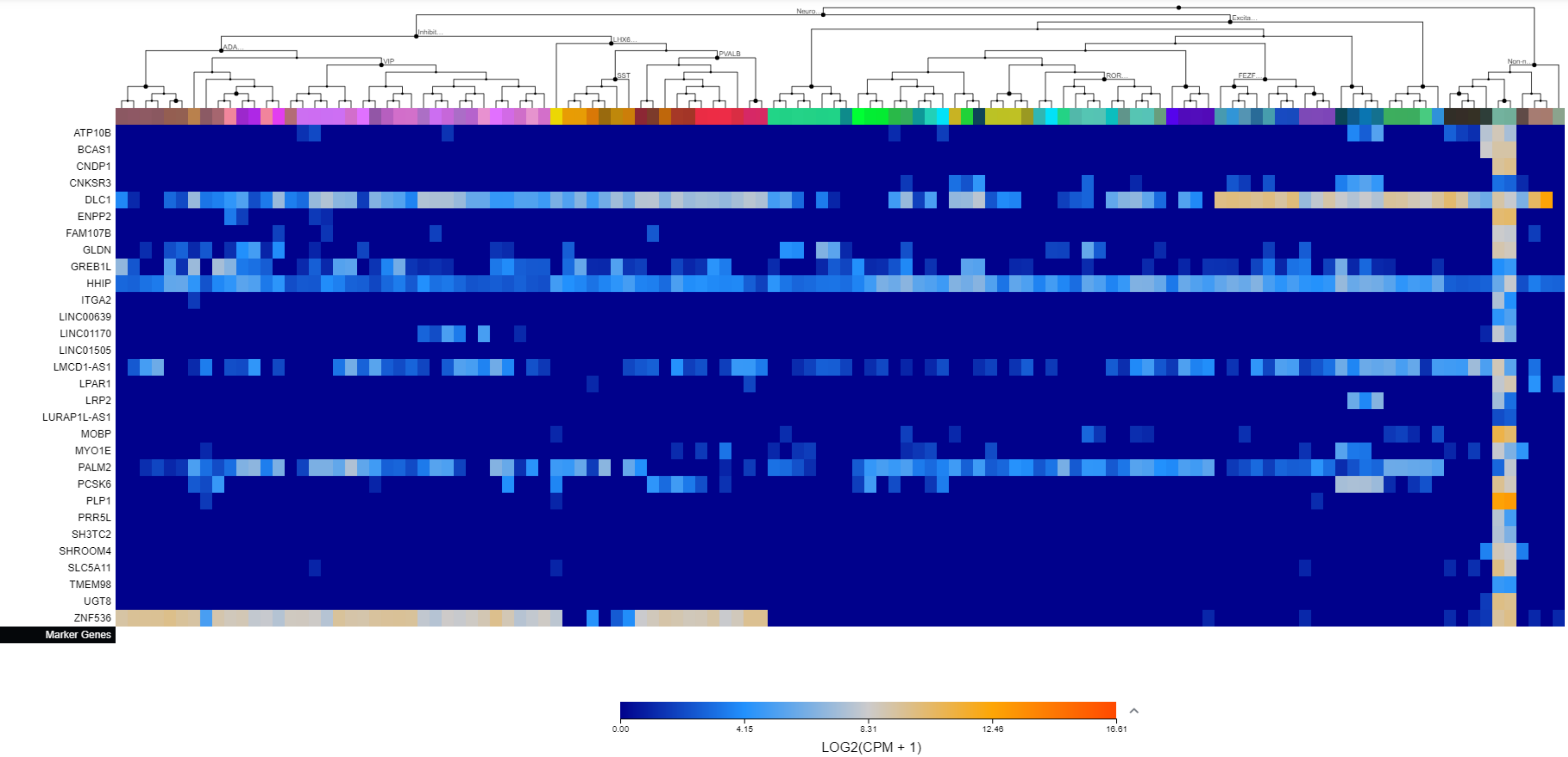

Supplementary File 3. The Allen Brain Map cell type heatmaps by cluster.

Cluster 8

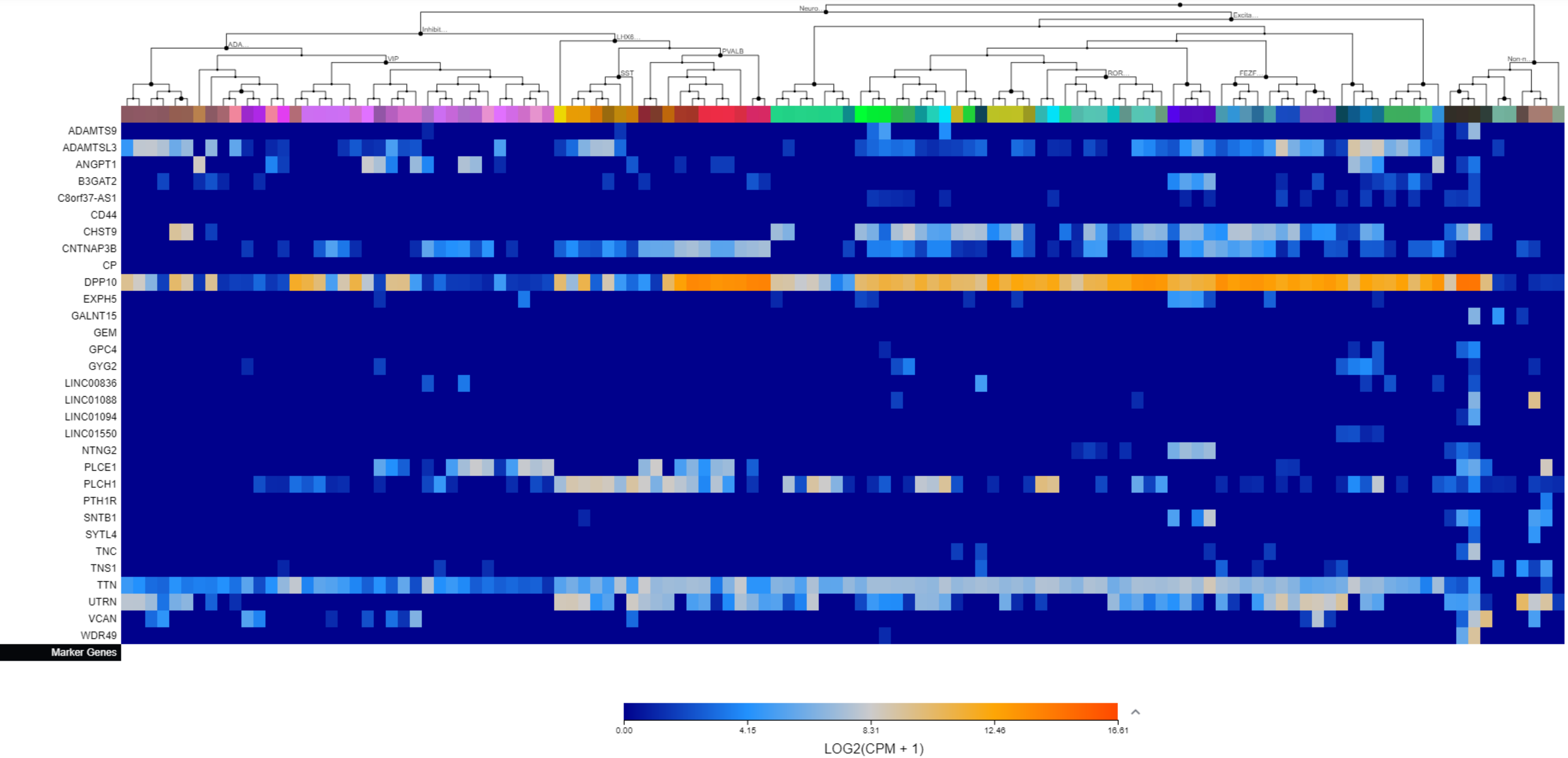

Supplementary File 3. The Allen Brain Map cell type heatmaps by cluster.

Cluster 9

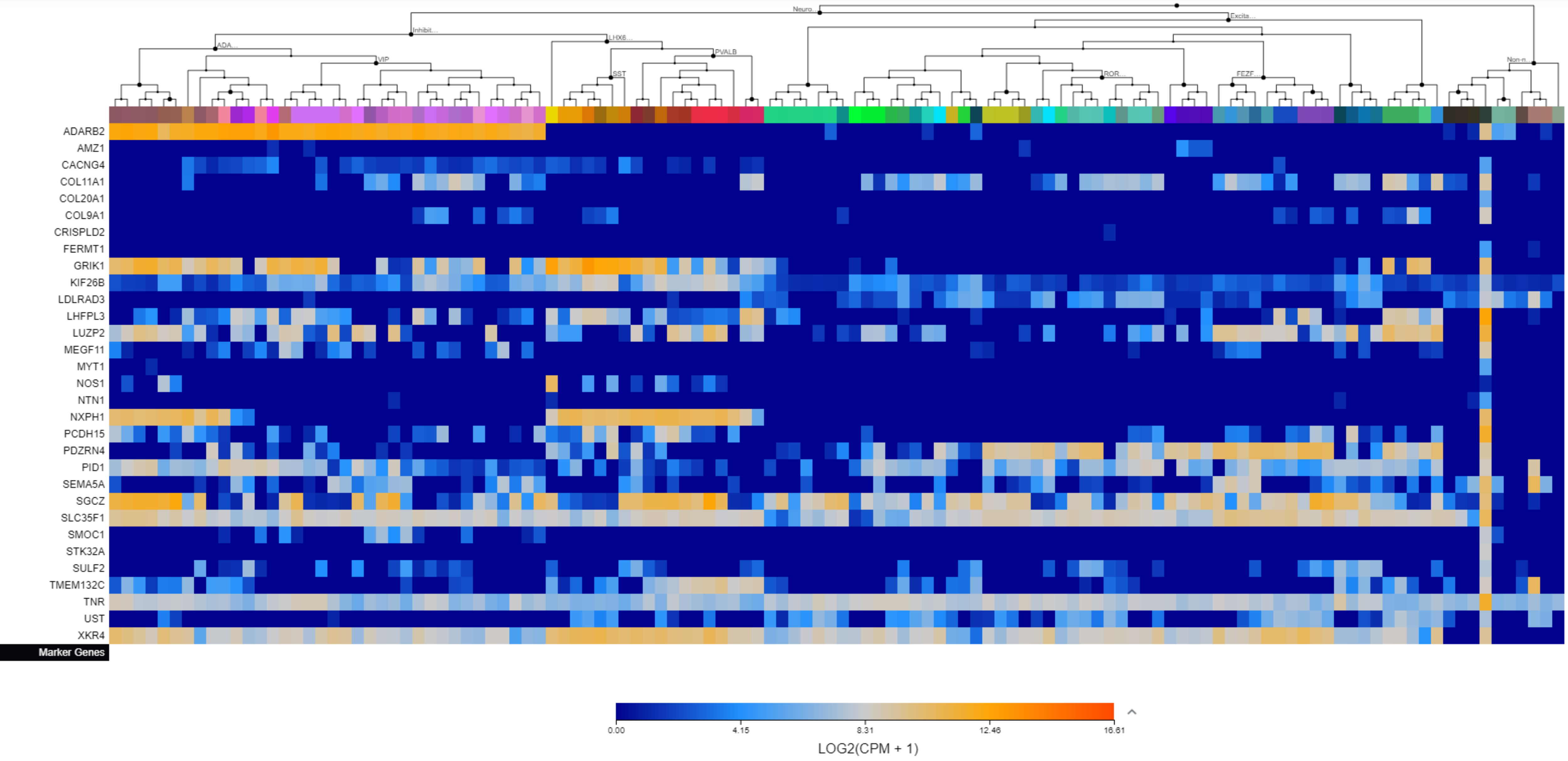

### Supplementary File 3. The Allen Brain Map cell type heatmaps by cluster.

#### Cluster 10

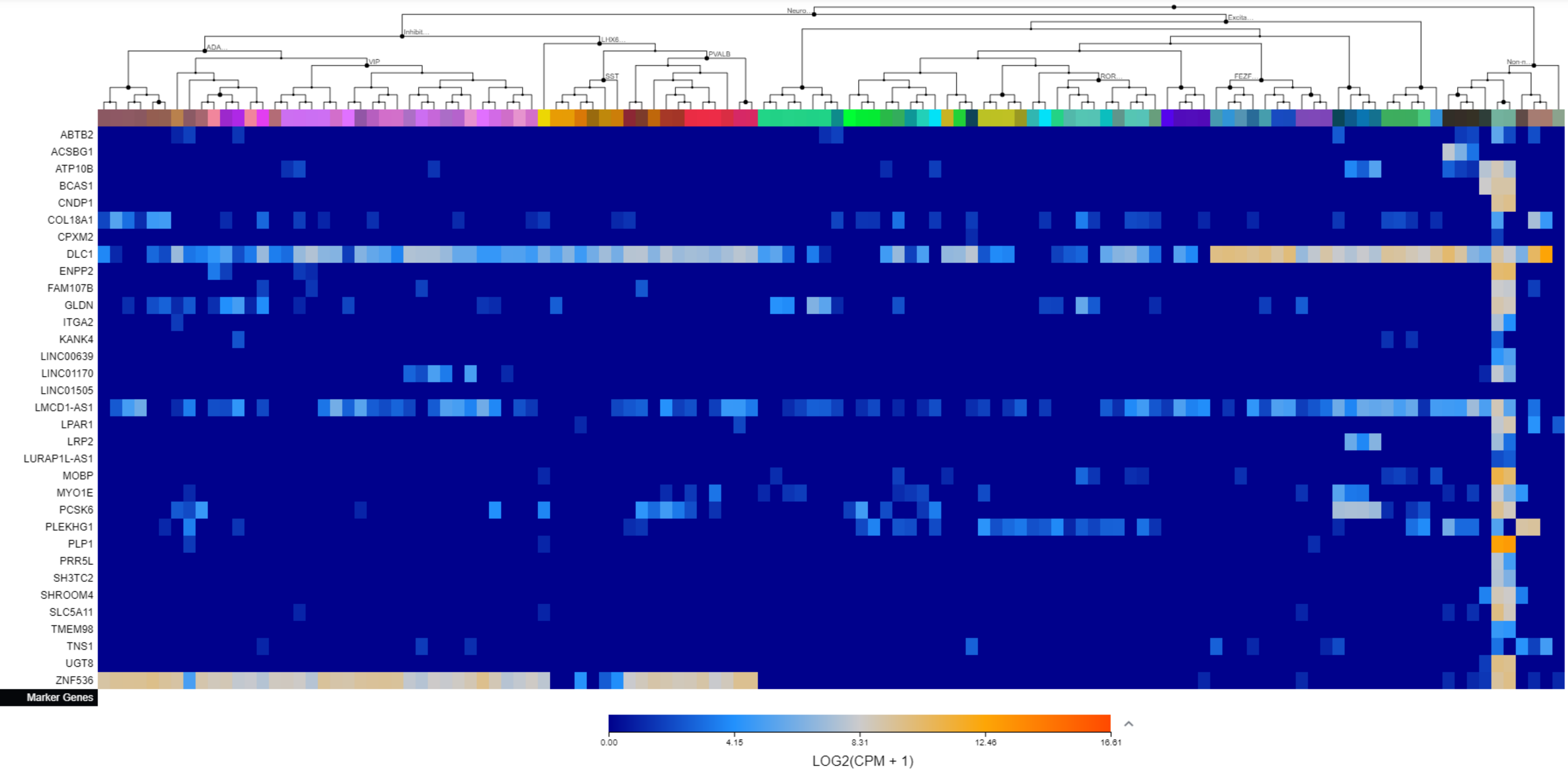

Supplementary File 3. The Allen Brain Map cell type heatmaps by cluster.

Cluster 11

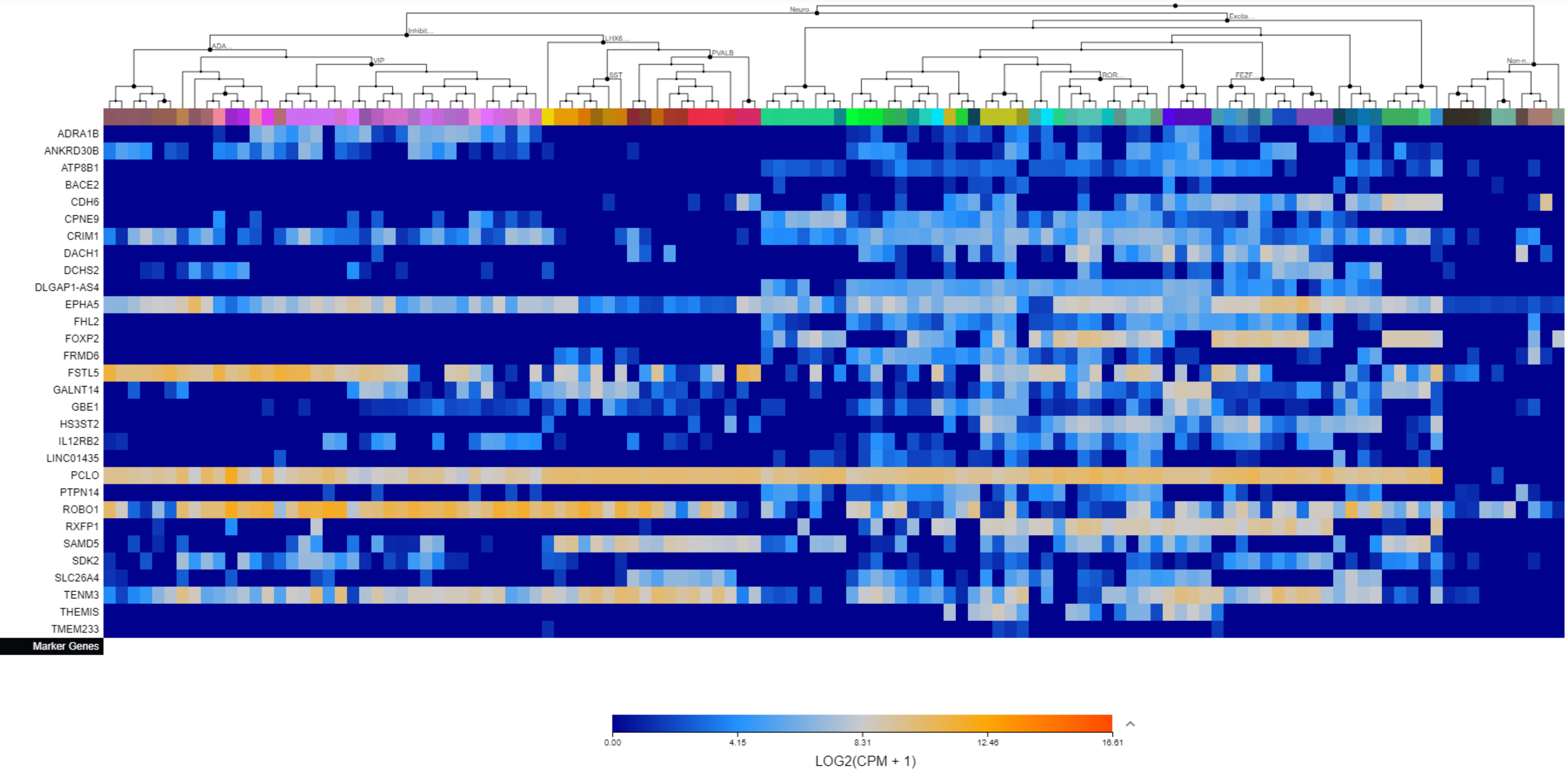

### Supplementary File 3. The Allen Brain Map cell type heatmaps by cluster.

#### Cluster 12

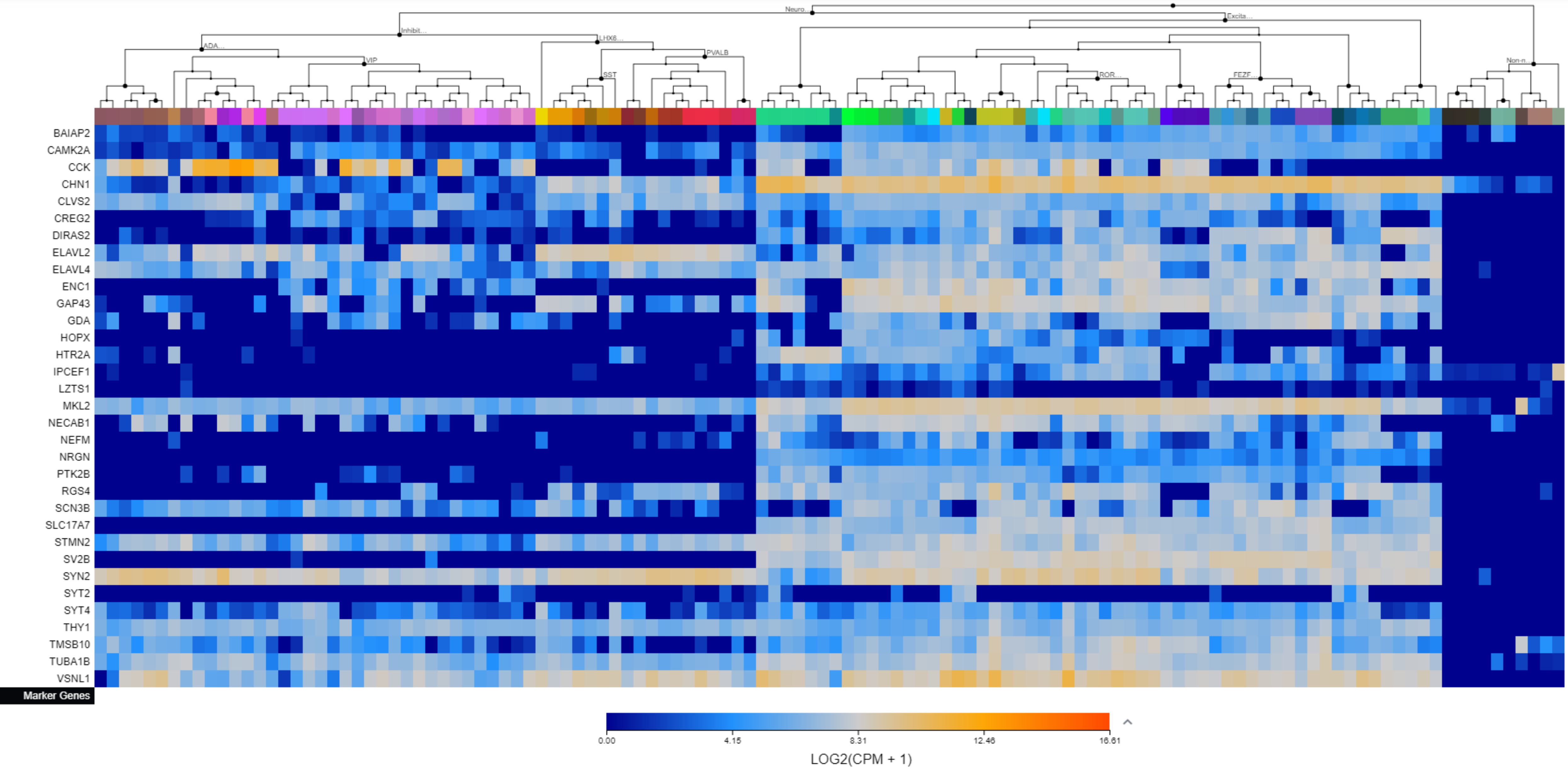

### Supplementary File 3. The Allen Brain Map cell type heatmaps by cluster.

#### Cluster 13

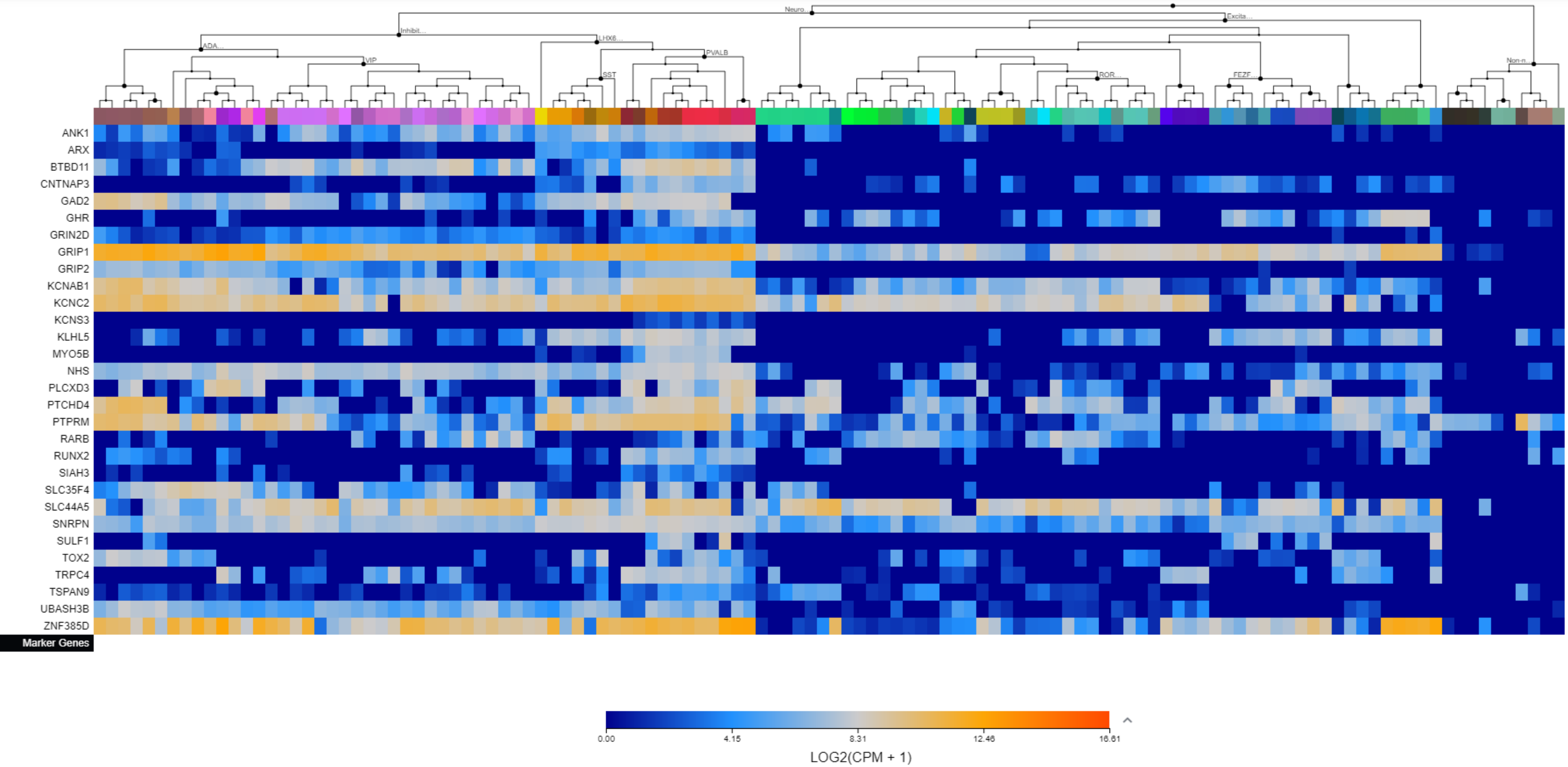

### Supplementary File 3. The Allen Brain Map cell type heatmaps by cluster.

Cluster 14

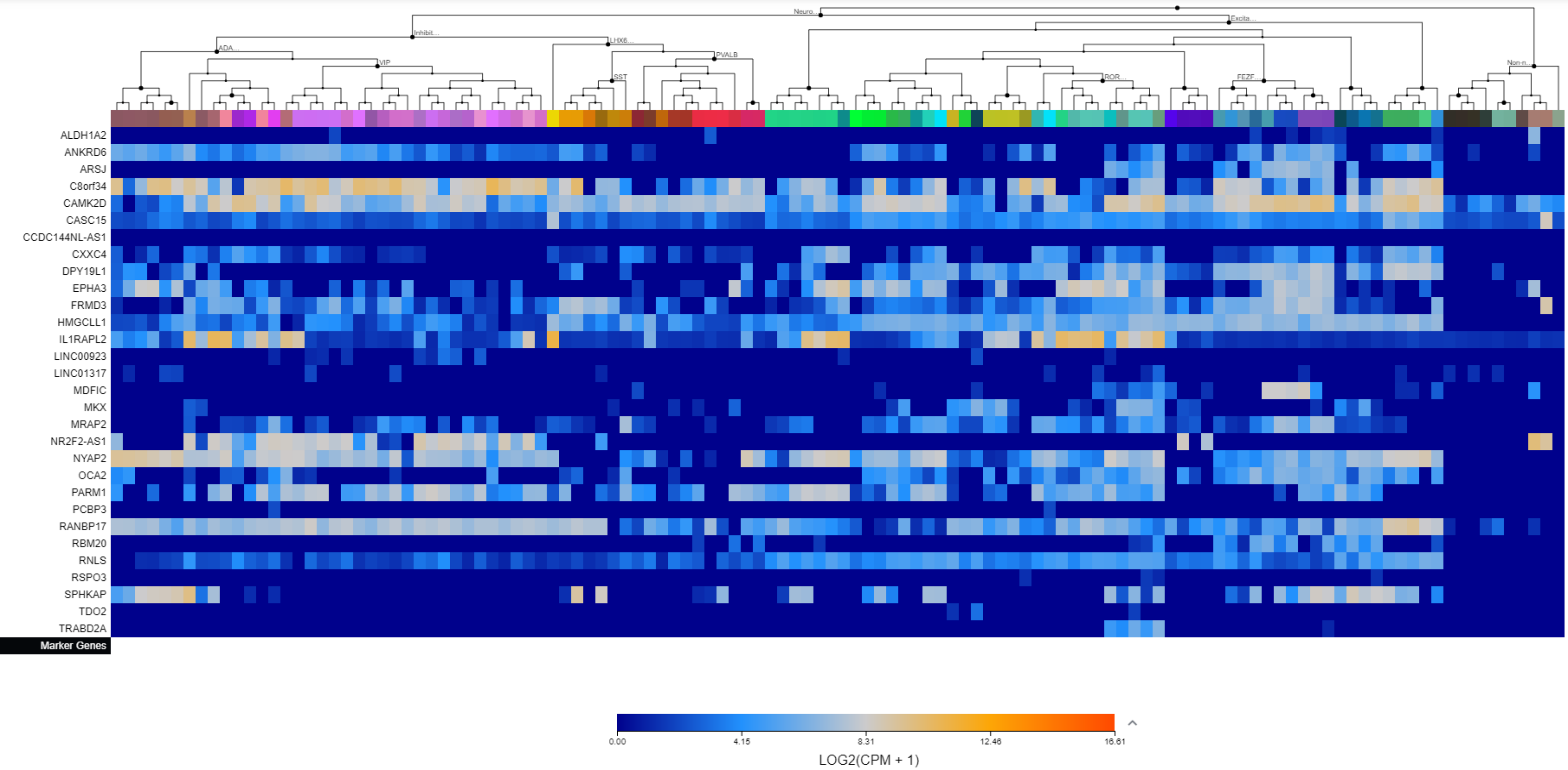

Supplementary File 3. The Allen Brain Map cell type heatmaps by cluster.

Cluster 15

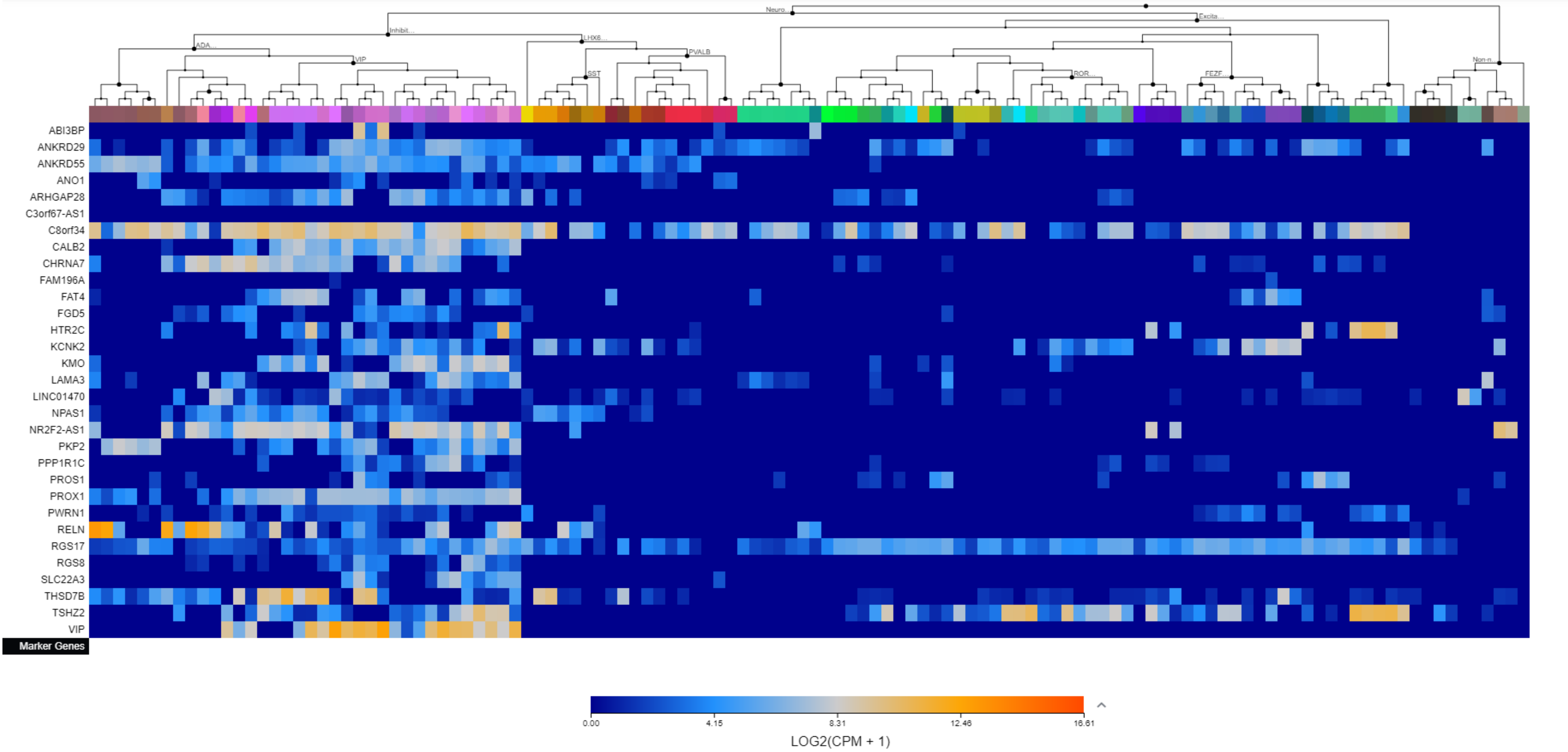

Supplementary File 3. The Allen Brain Map cell type heatmaps by cluster.

Cluster 16

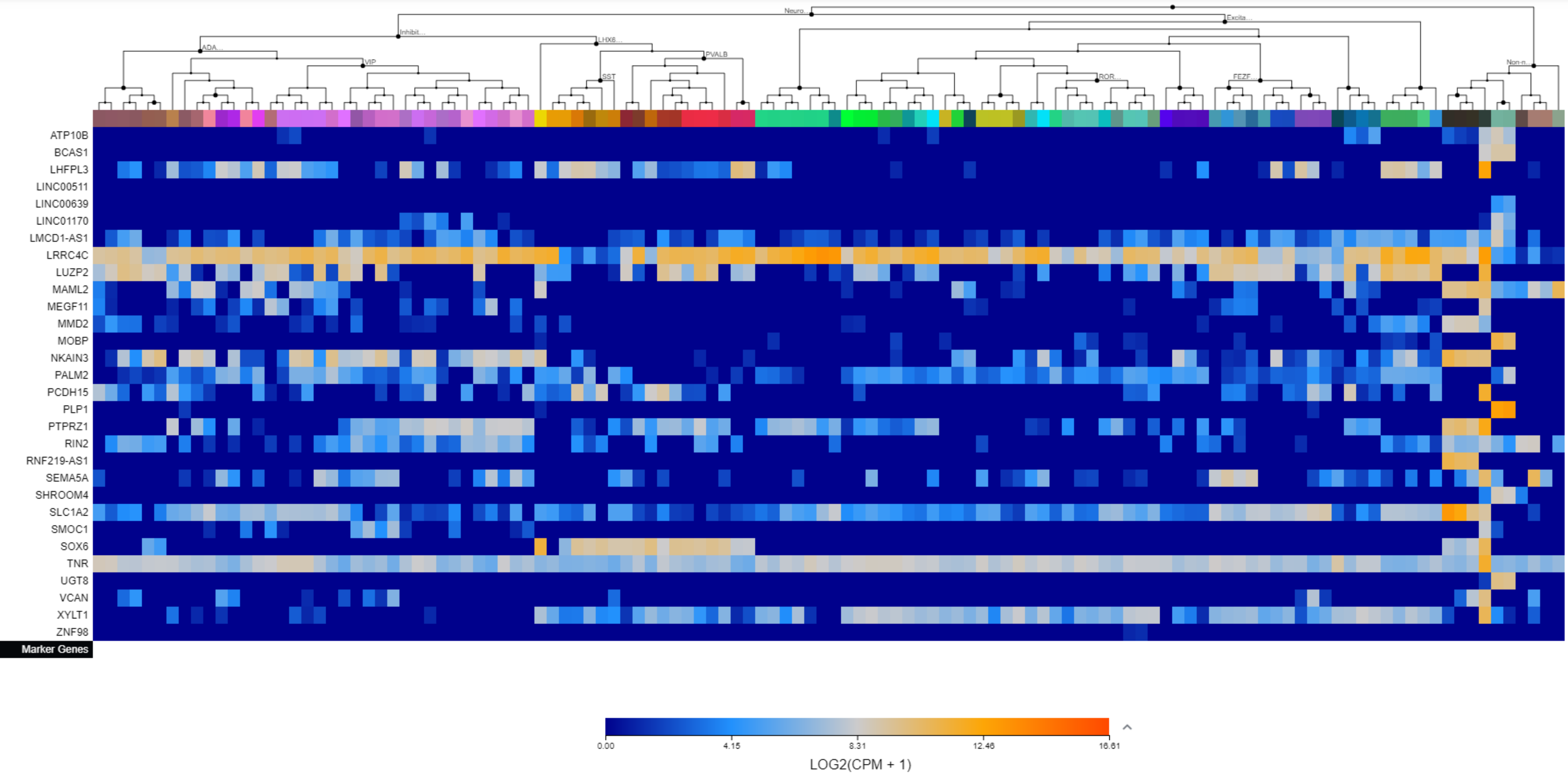

Supplementary File 3. The Allen Brain Map cell type heatmaps by cluster.

Cluster 17

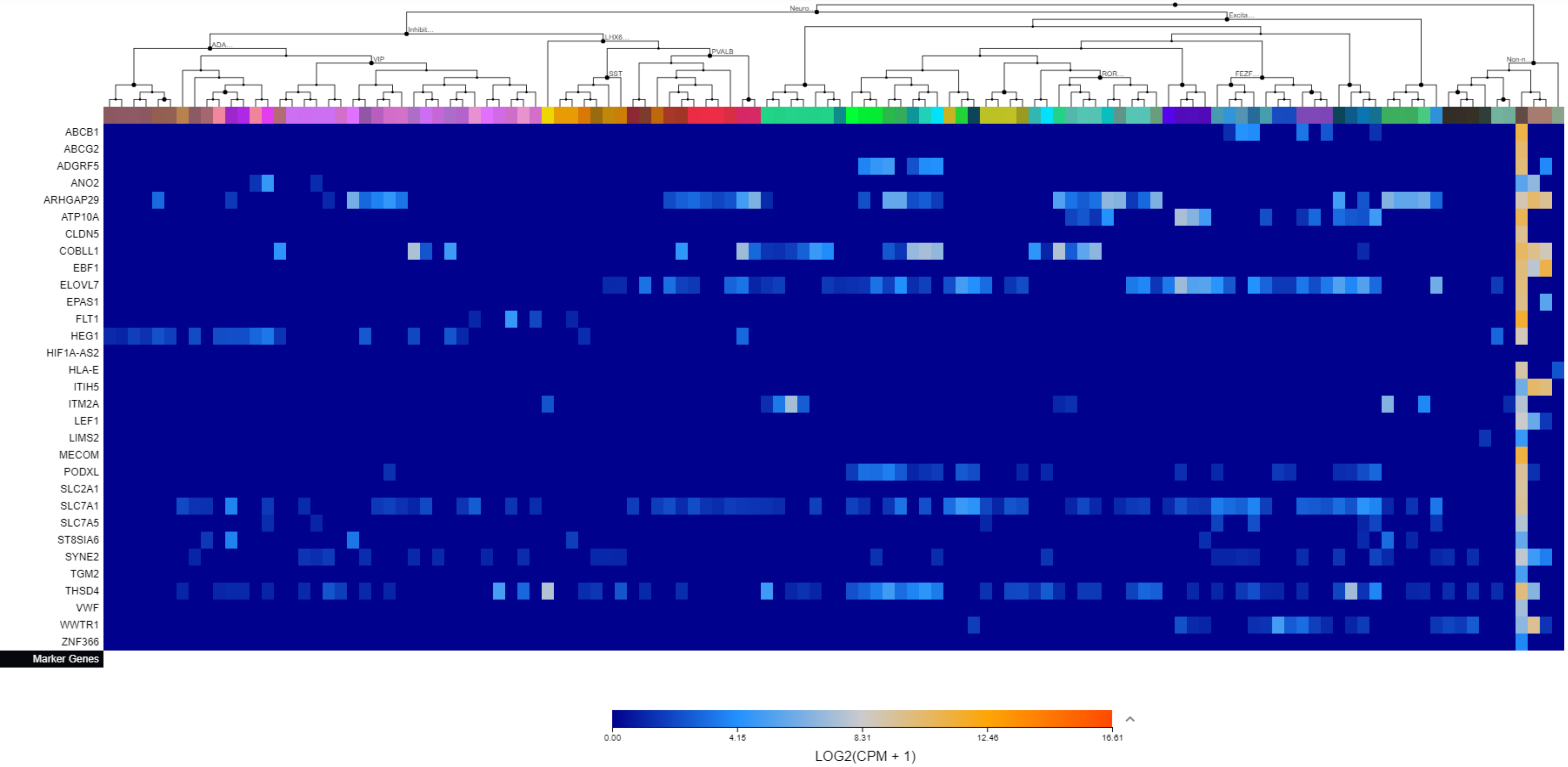

Supplementary File 3. The Allen Brain Map cell type heatmaps by cluster.

Cluster 18

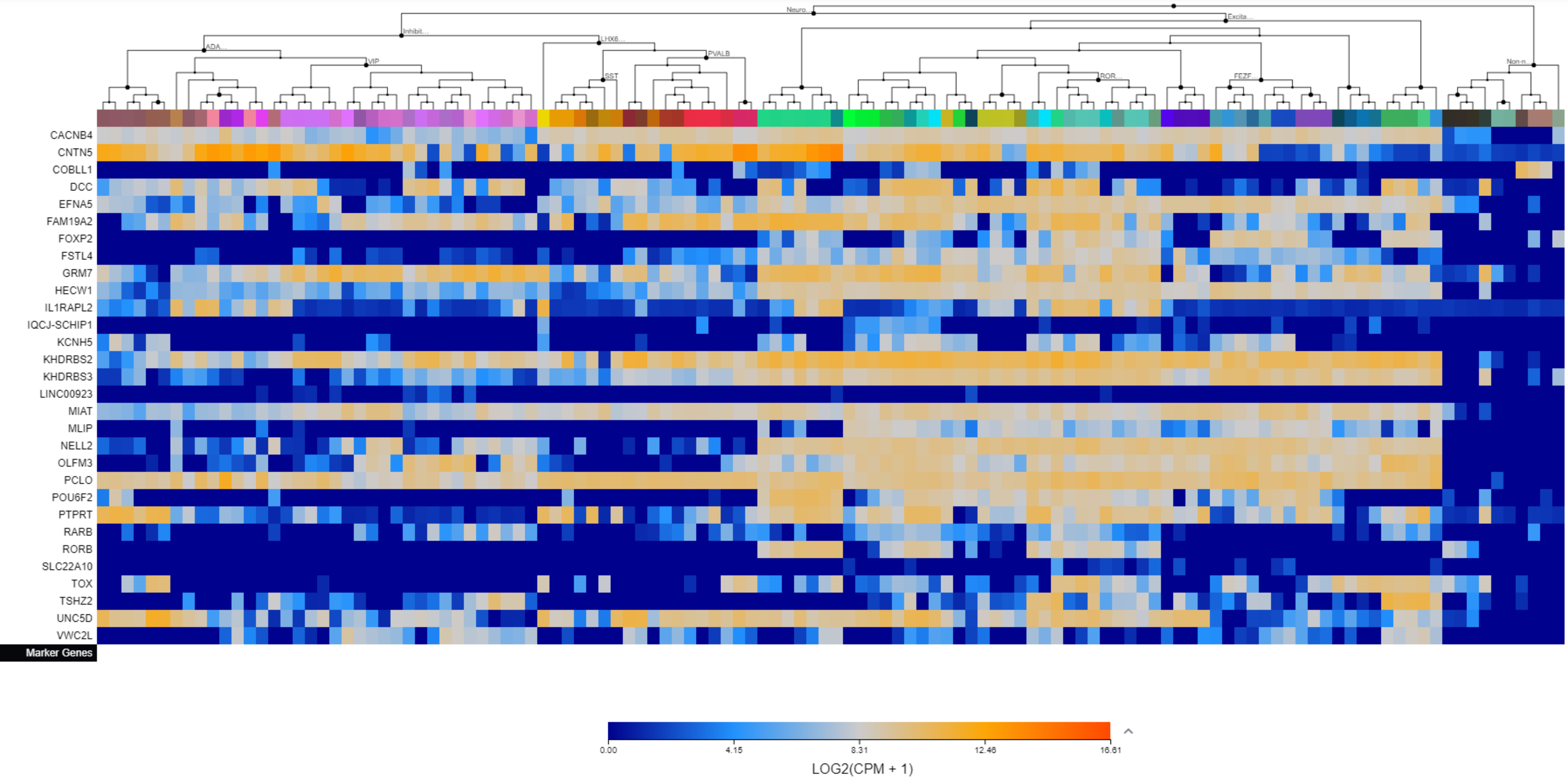

Supplementary File 3. The Allen Brain Map cell type heatmaps by cluster.

Cluster 19

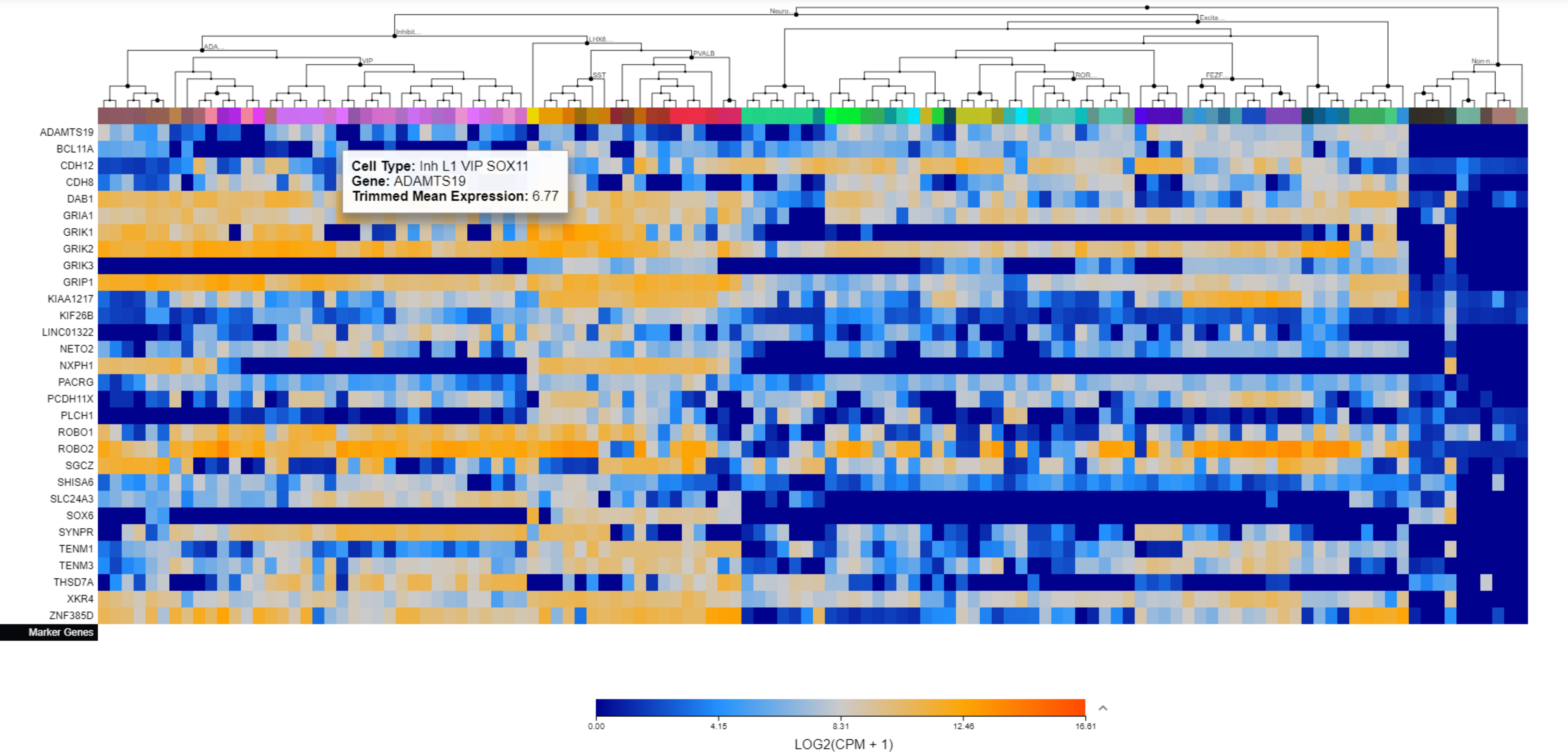

Supplementary File 3. The Allen Brain Map cell type heatmaps by cluster.

Cluster 20

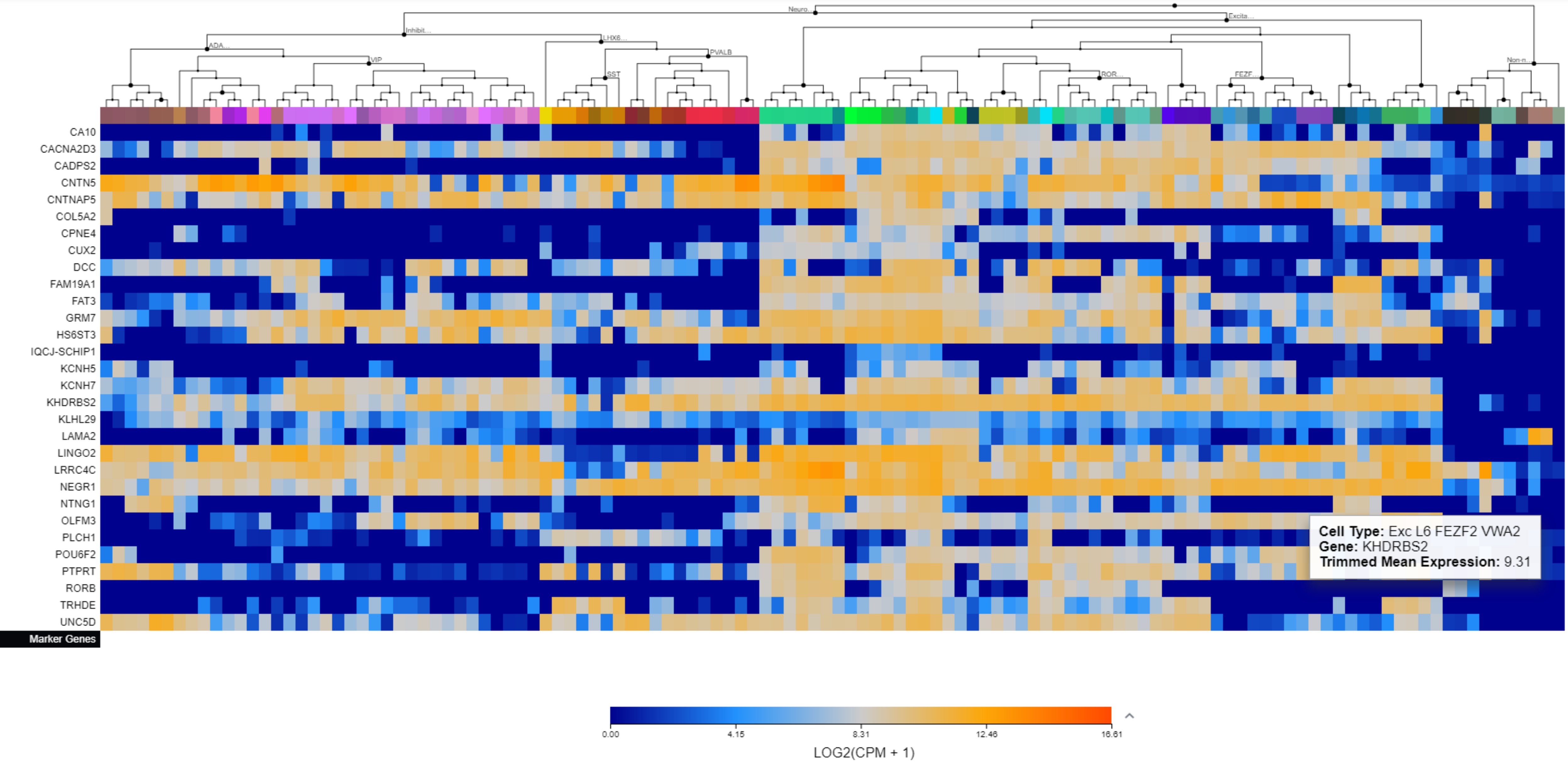

Supplementary File 3. The Allen Brain Map cell type heatmaps by cluster.

Cluster 21

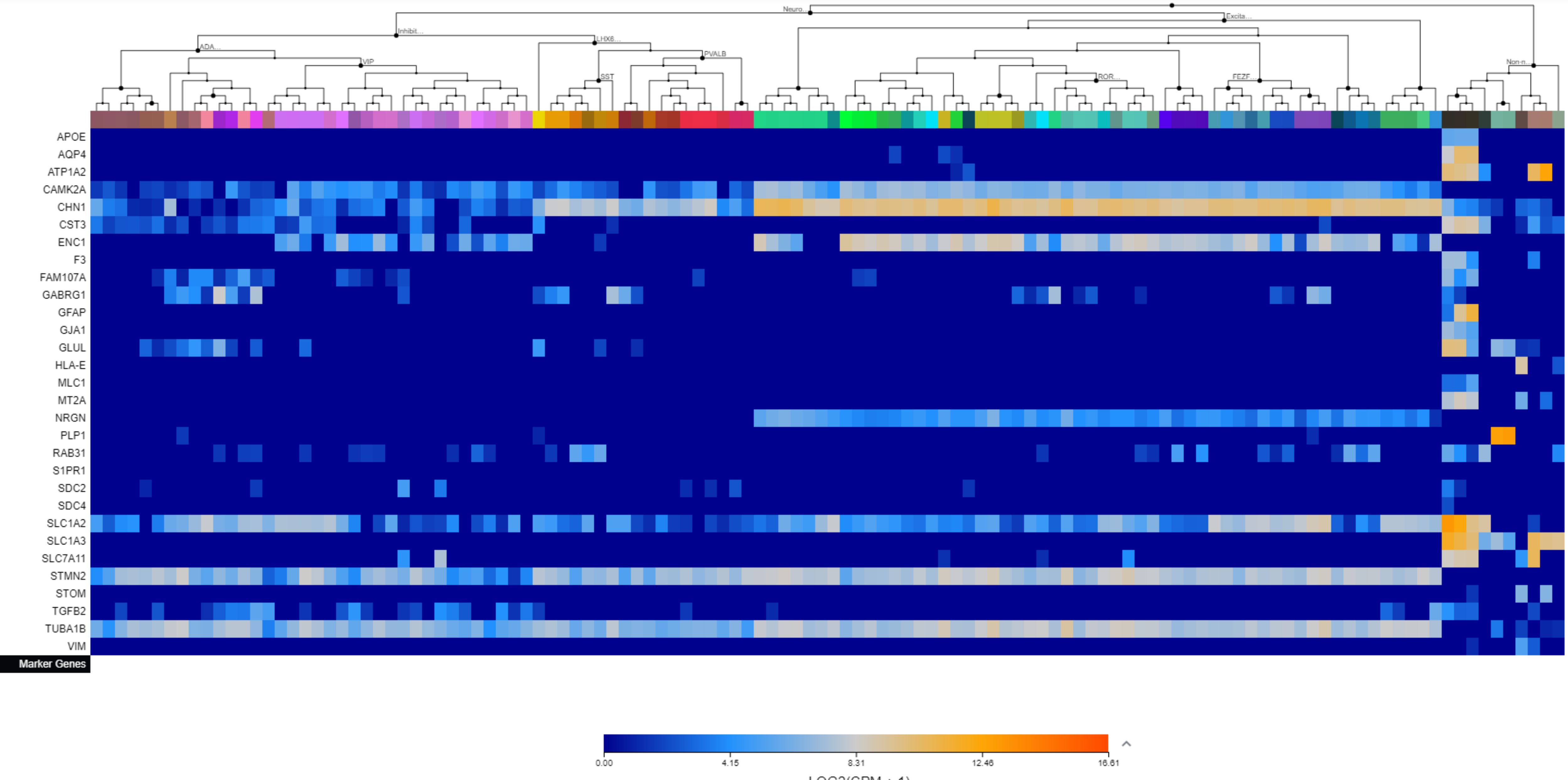

Supplementary File 3. The Allen Brain Map cell type heatmaps by cluster.

Cluster 22

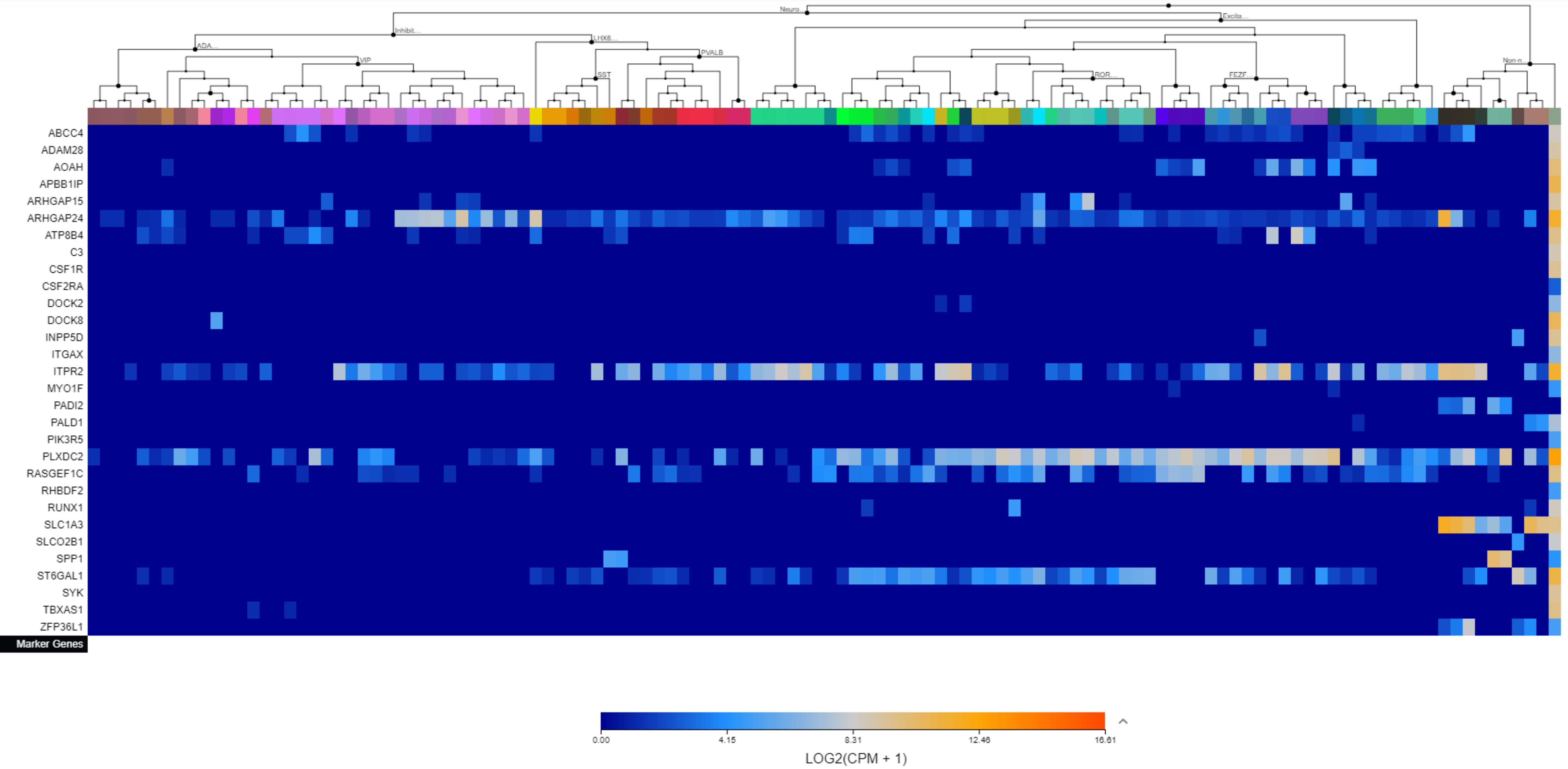

Supplementary File 3. The Allen Brain Map cell type heatmaps by cluster.

Cluster 23

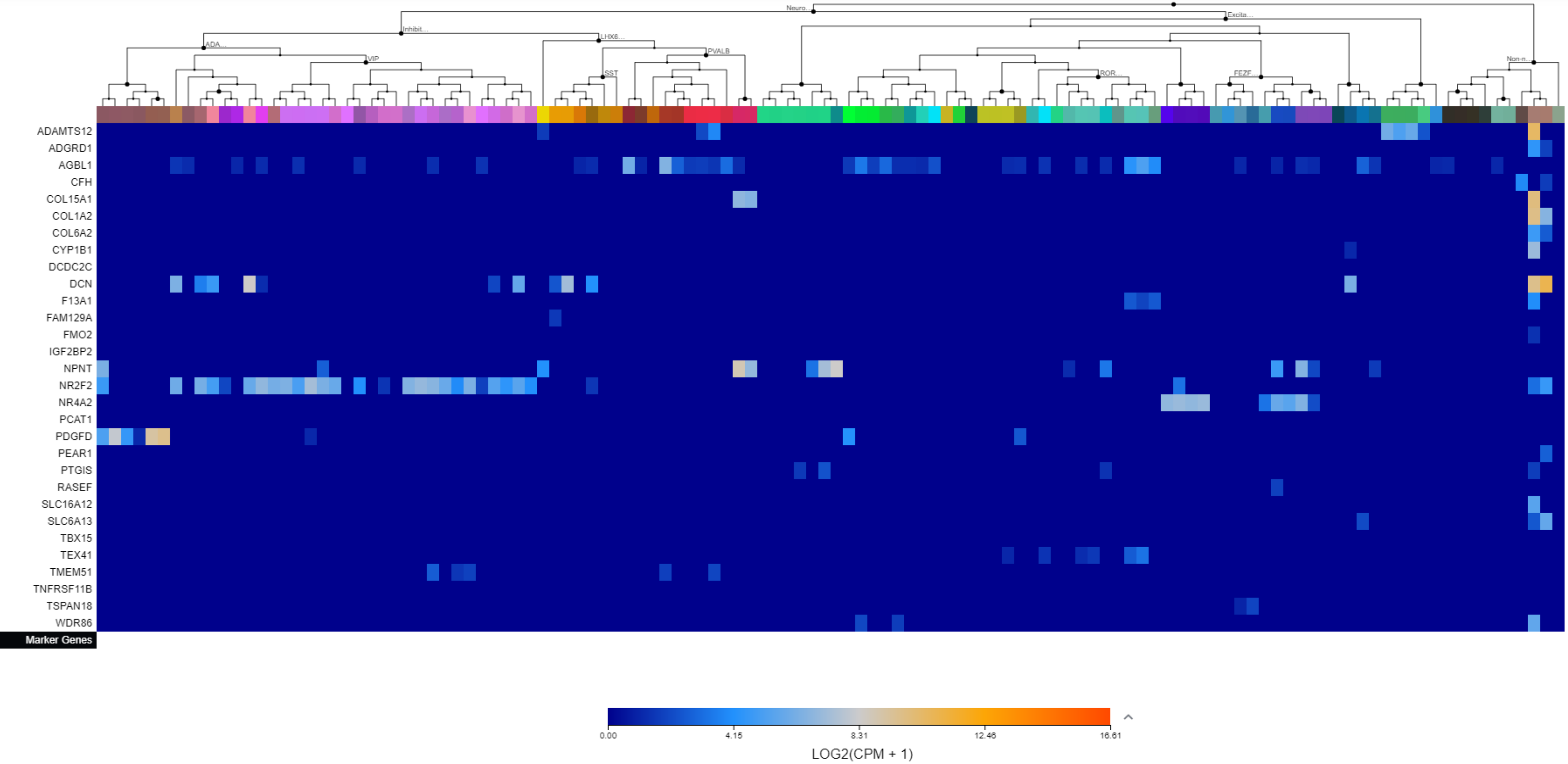

### Supplementary File 3. The Allen Brain Map cell type heatmaps by cluster.

#### Cluster 24

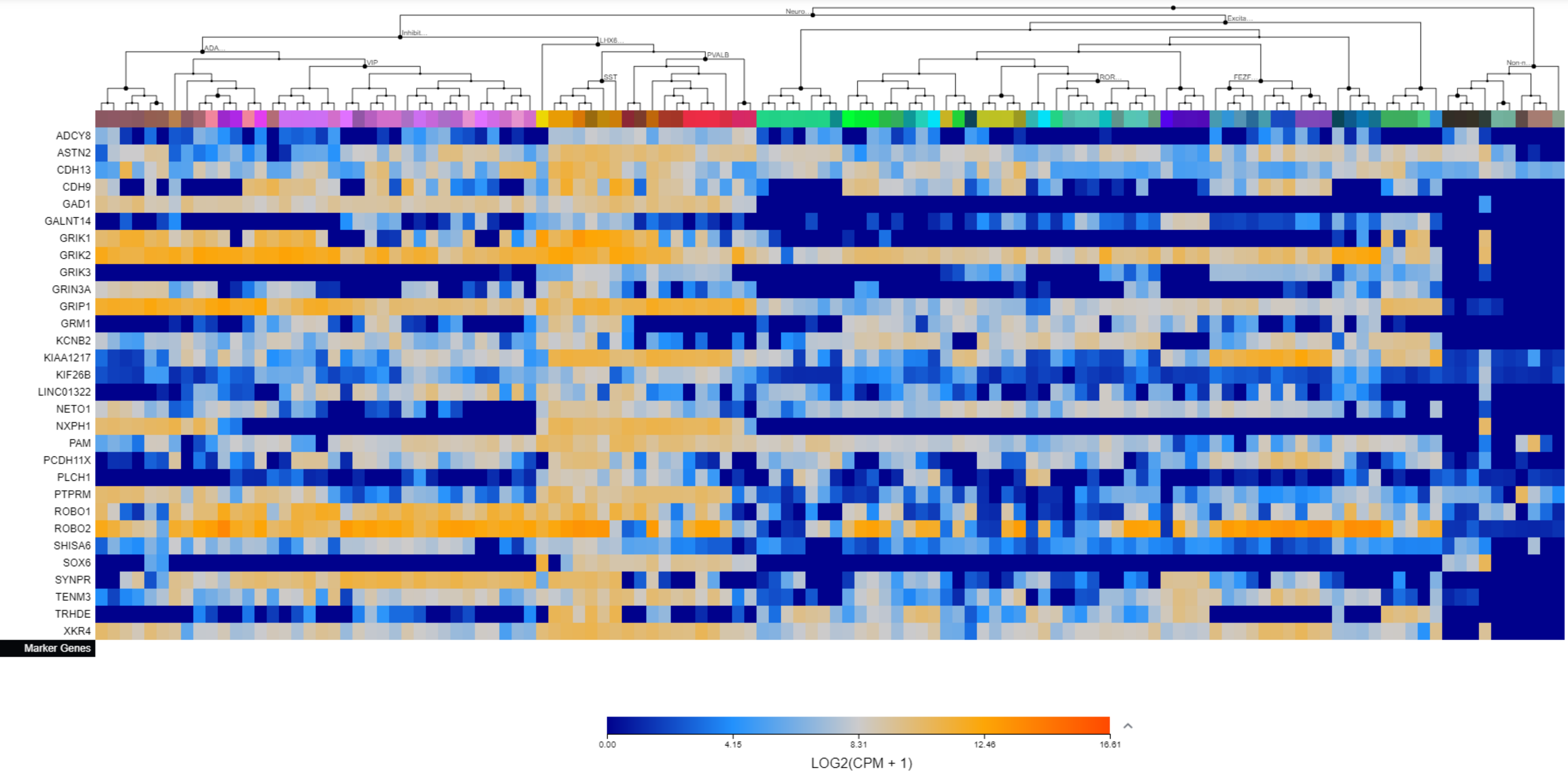

Supplementary File 3. The Allen Brain Map cell type heatmaps by cluster.

Cluster 25

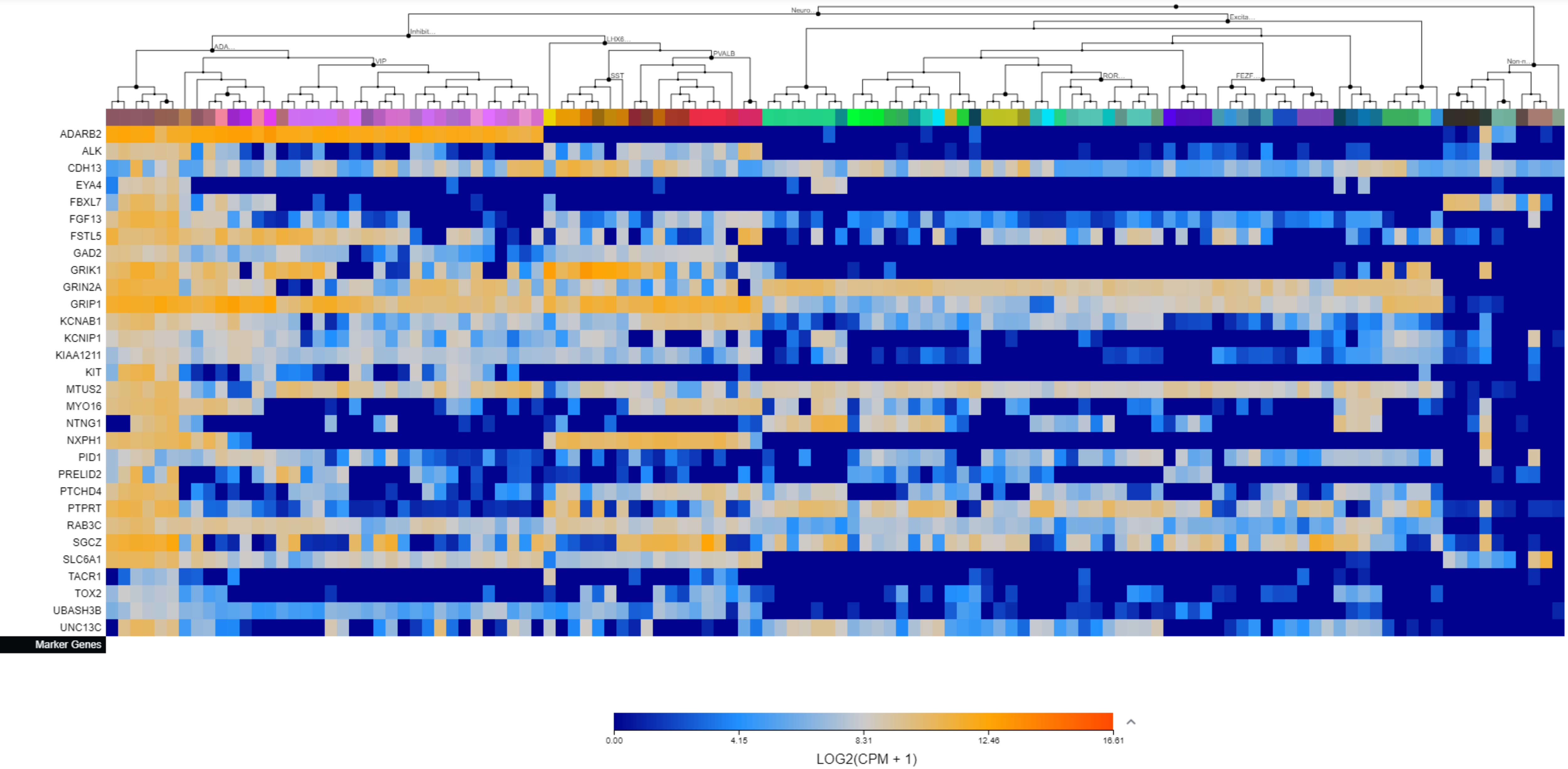

Supplementary File 3. The Allen Brain Map cell type heatmaps by cluster.

Cluster 26

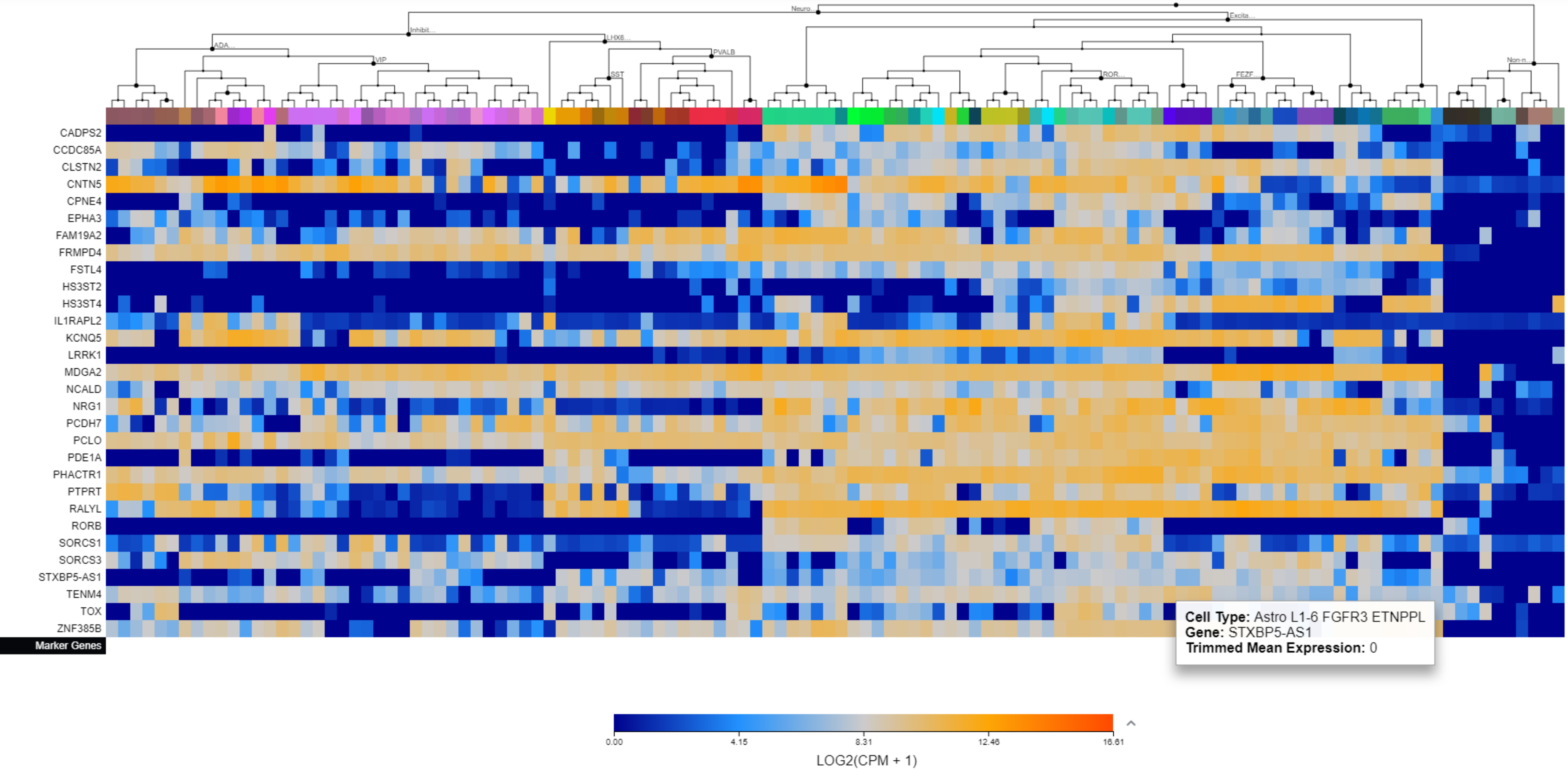

Supplementary File 3. The Allen Brain Map cell type heatmaps by cluster.

Cluster 27

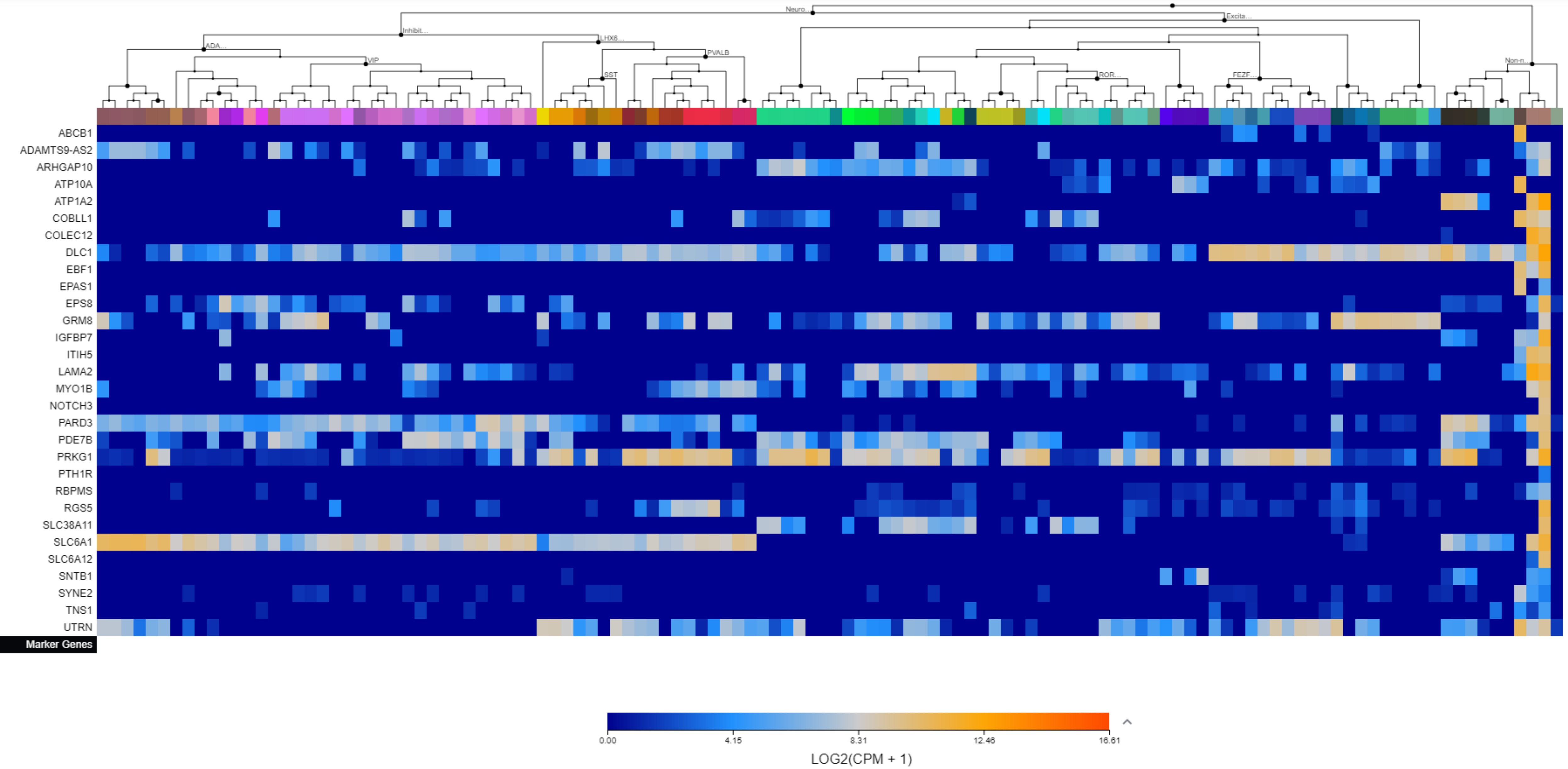

Supplementary File 3. The Allen Brain Map cell type heatmaps by cluster.

Cluster 28

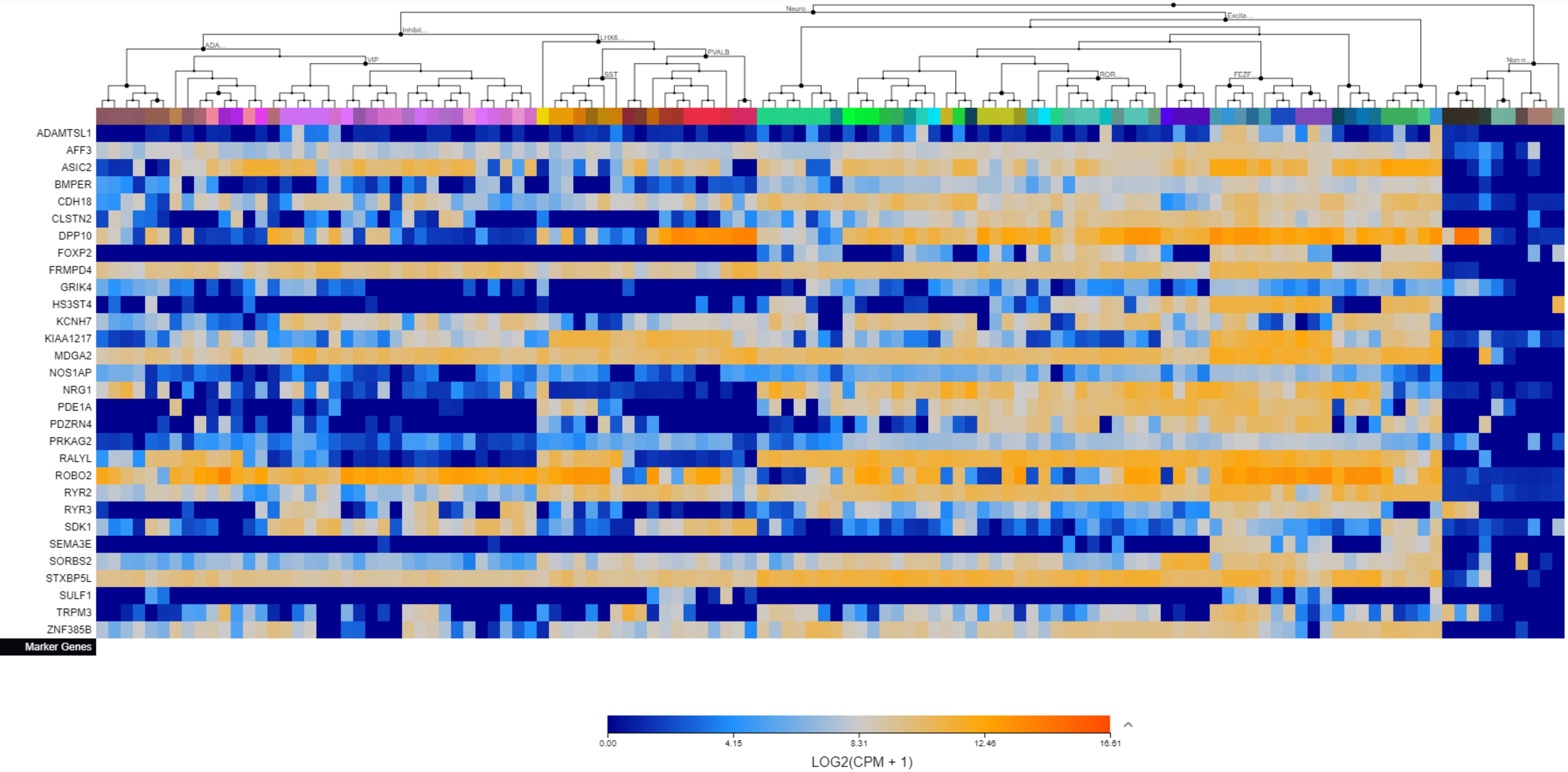

### Supplementary File 3. The Allen Brain Map cell type heatmaps by cluster.

#### Cluster 29

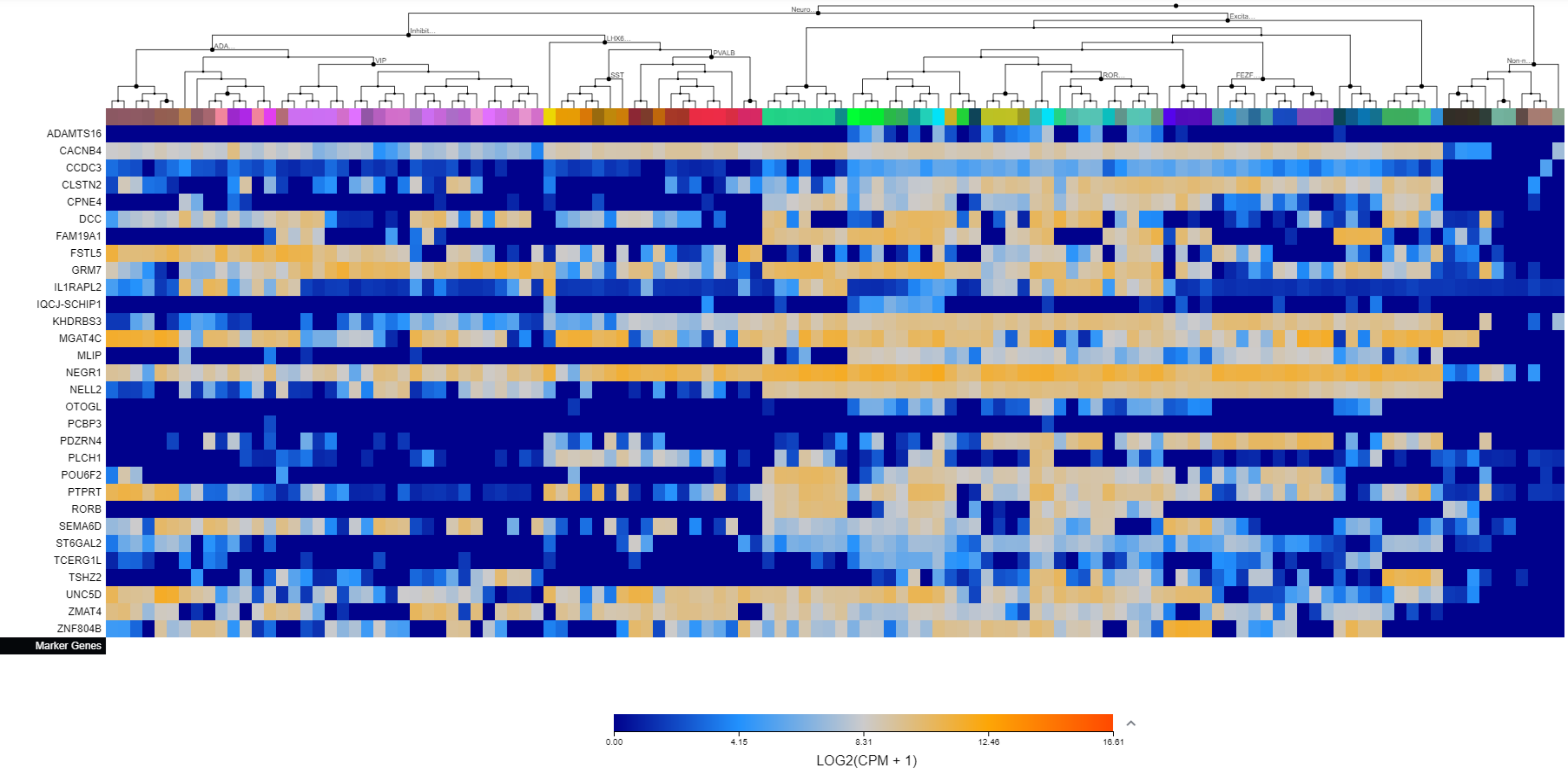

Supplementary File 3. The Allen Brain Map cell type heatmaps by cluster.

Cluster 30

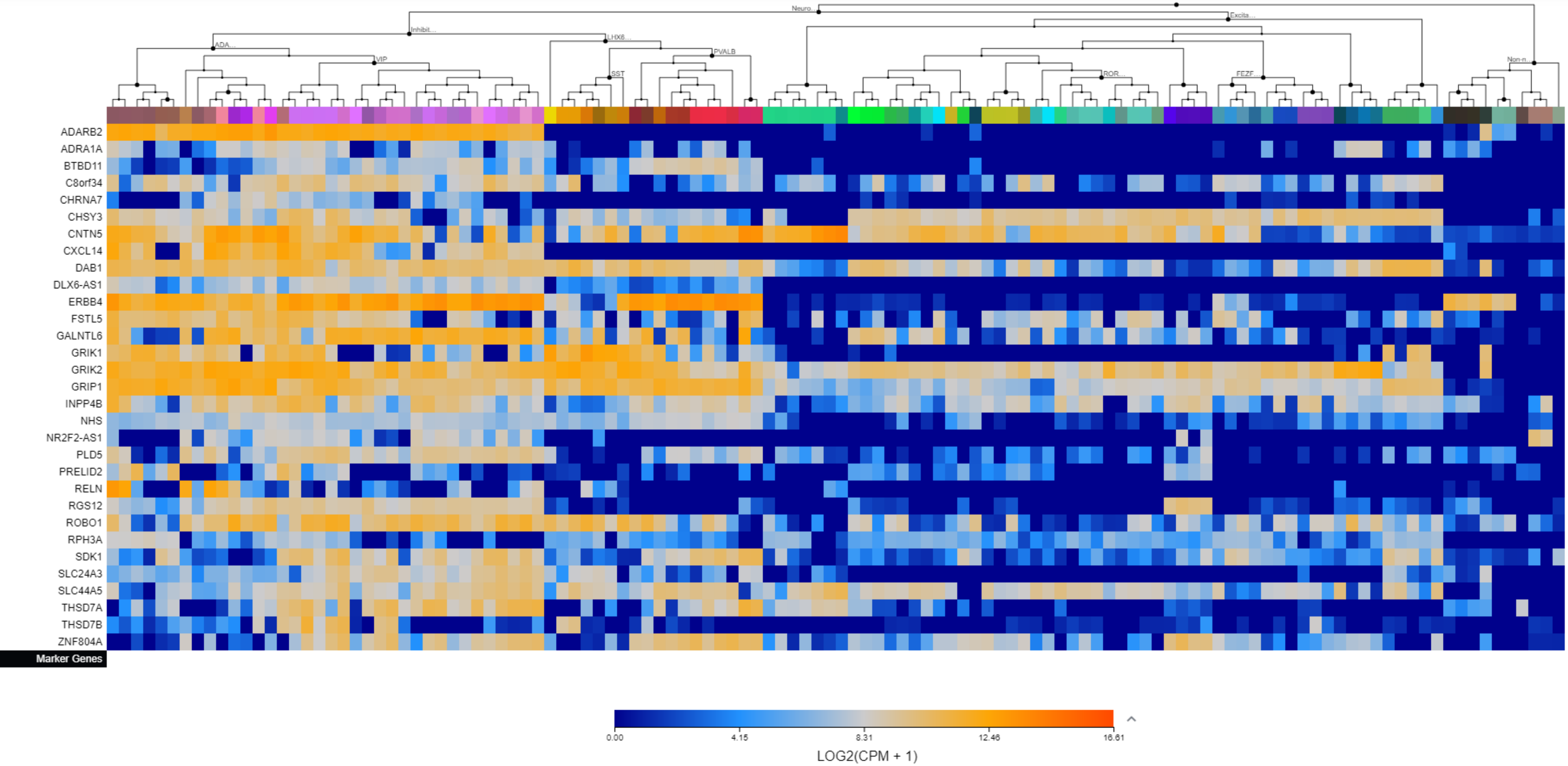

Supplementary File 3. The Allen Brain Map cell type heatmaps by cluster.

Cluster 31

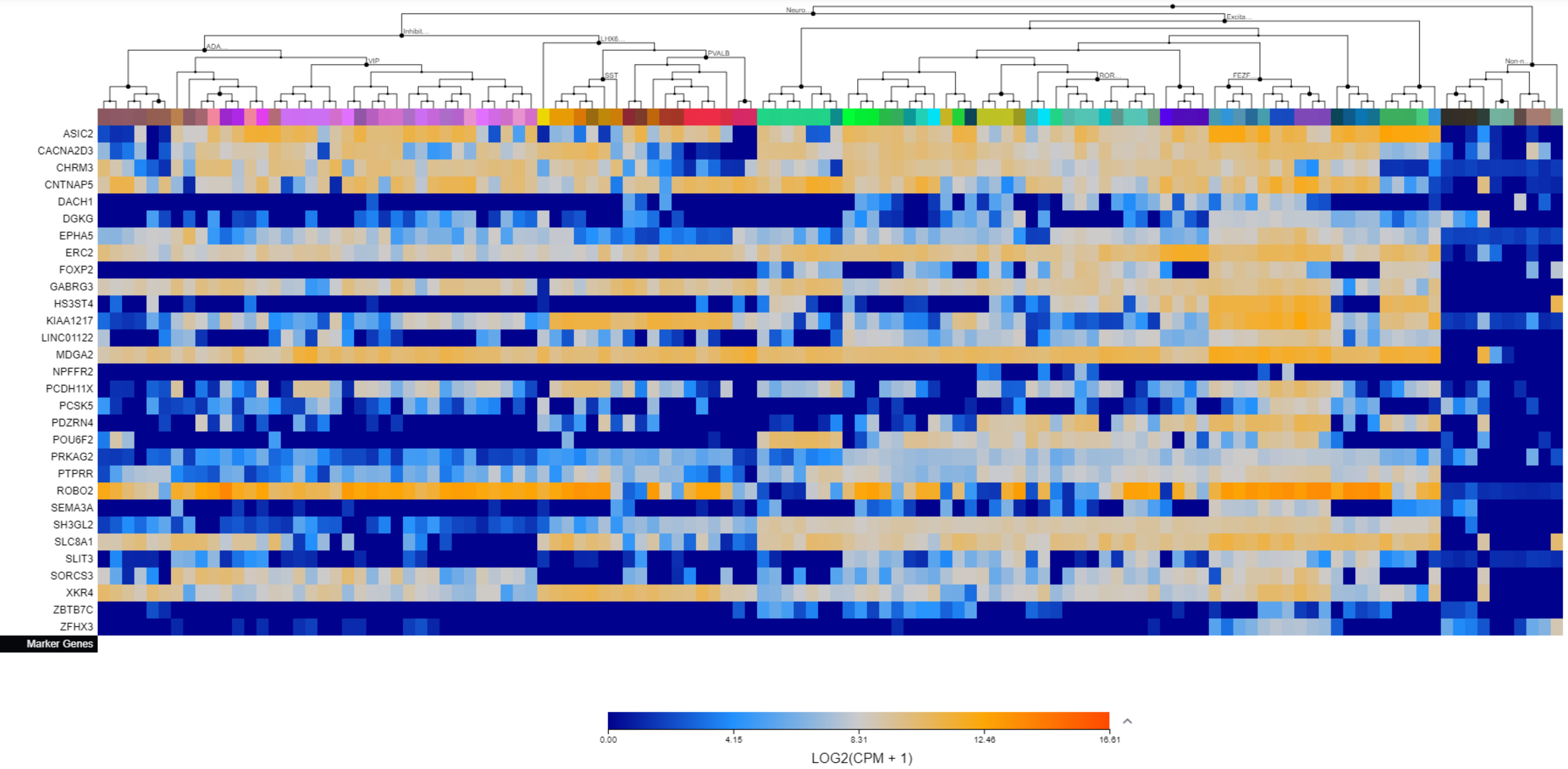

Supplementary File 3. The Allen Brain Map cell type heatmaps by cluster.

Cluster 32

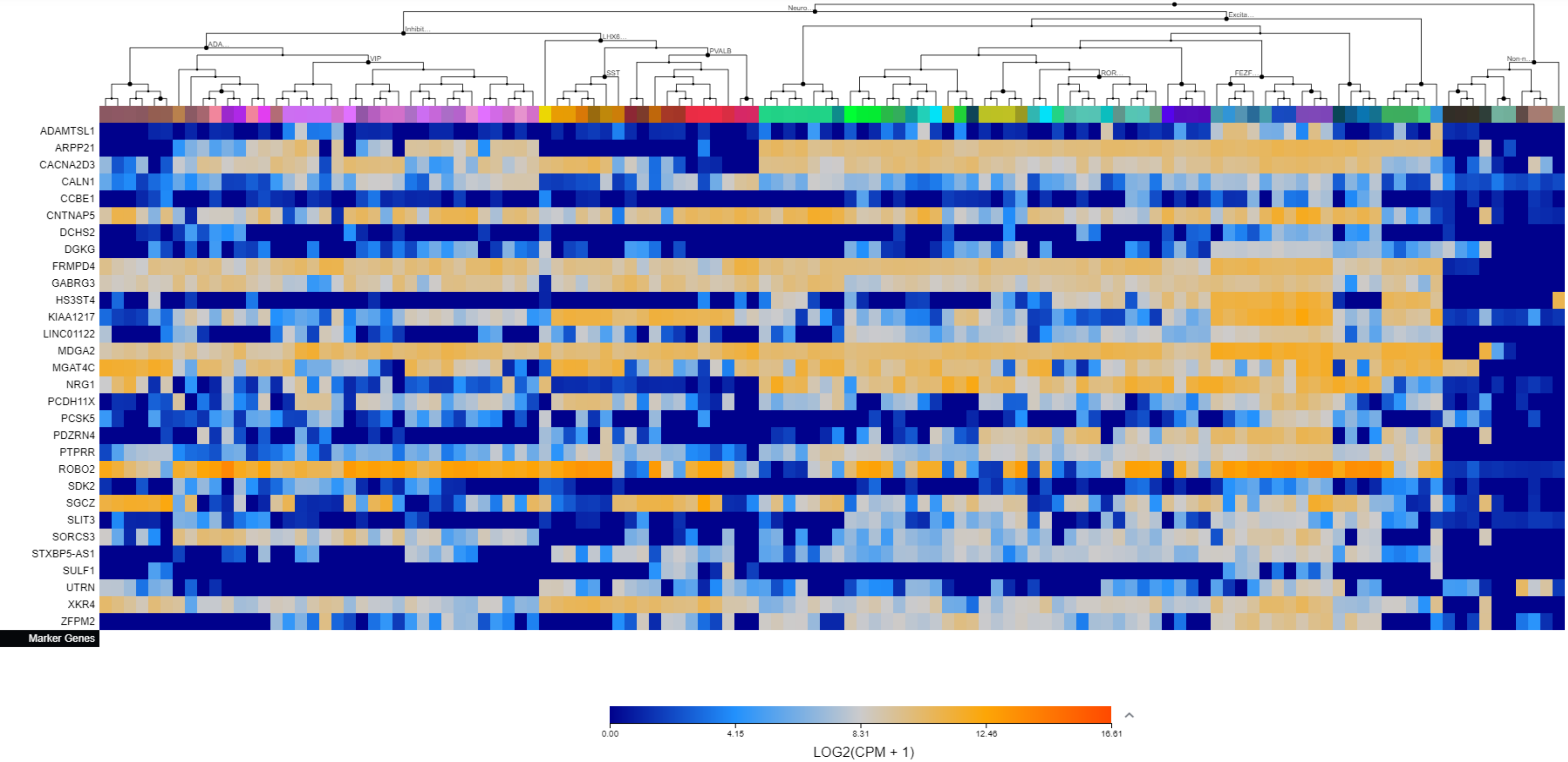

Supplementary File 3. The Allen Brain Map cell type heatmaps by cluster.

Cluster 33

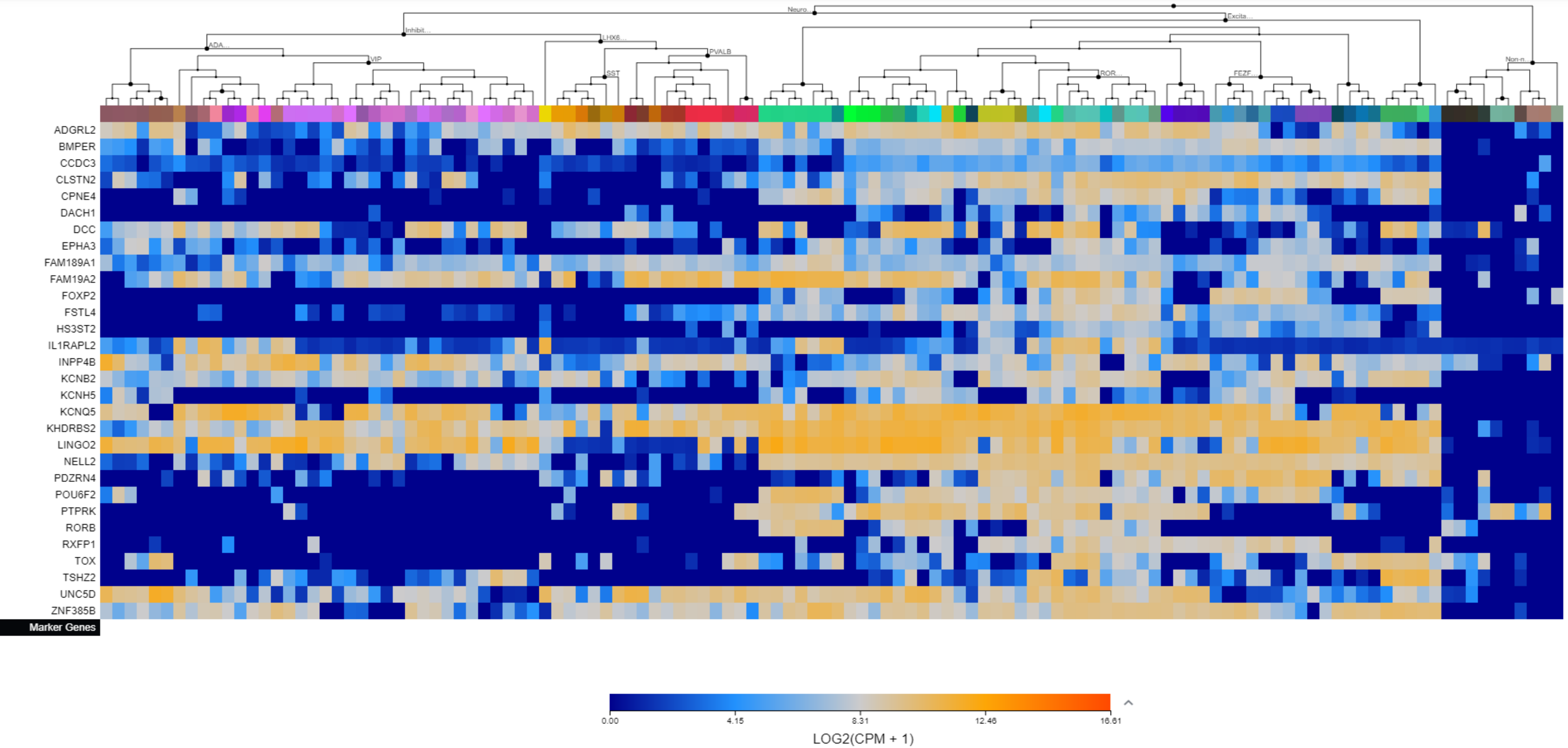

Supplementary File 3. The Allen Brain Map cell type heatmaps by cluster.

Cluster 34

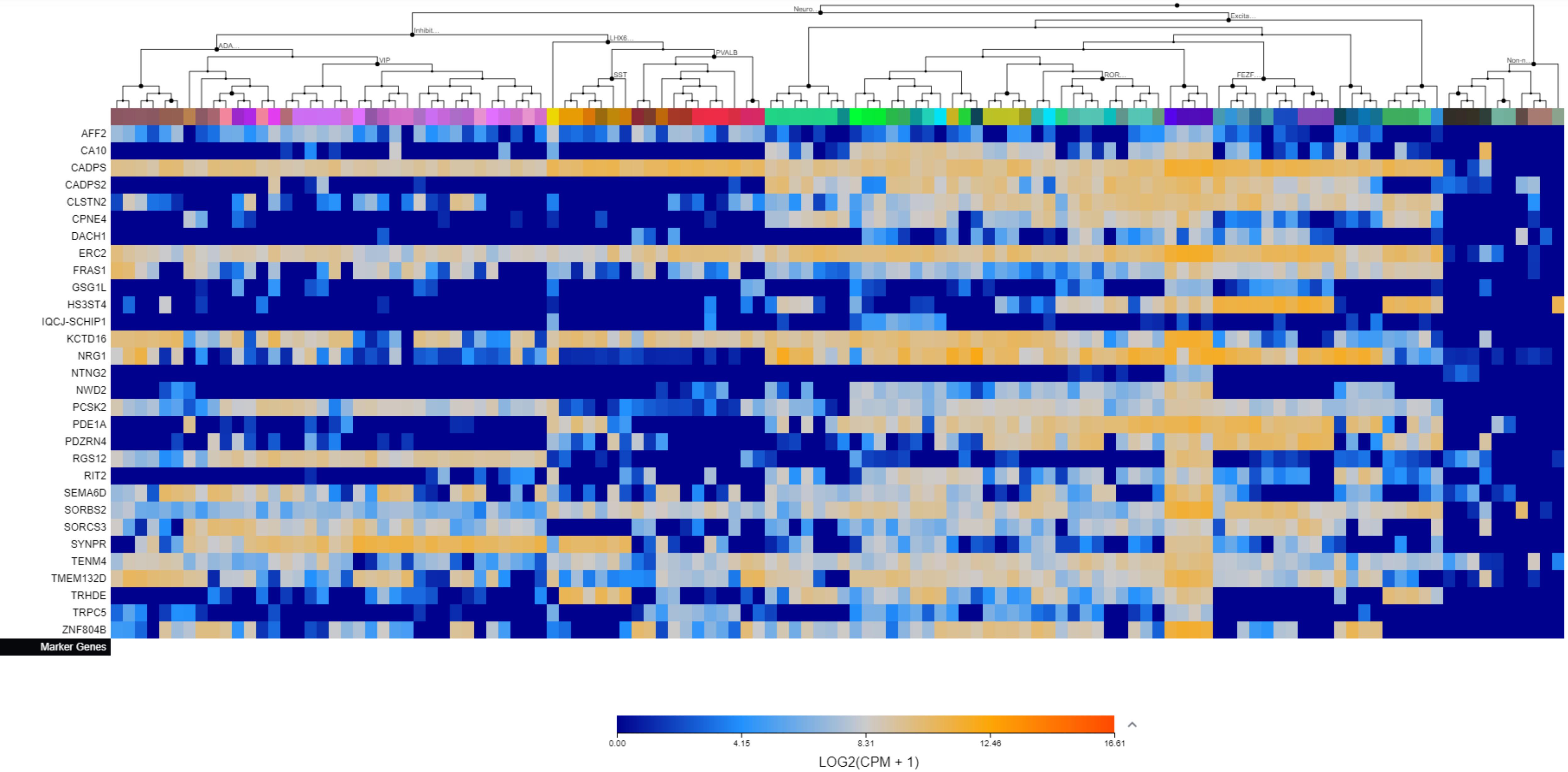

Supplementary File 3. The Allen Brain Map cell type heatmaps by cluster.

Cluster 35

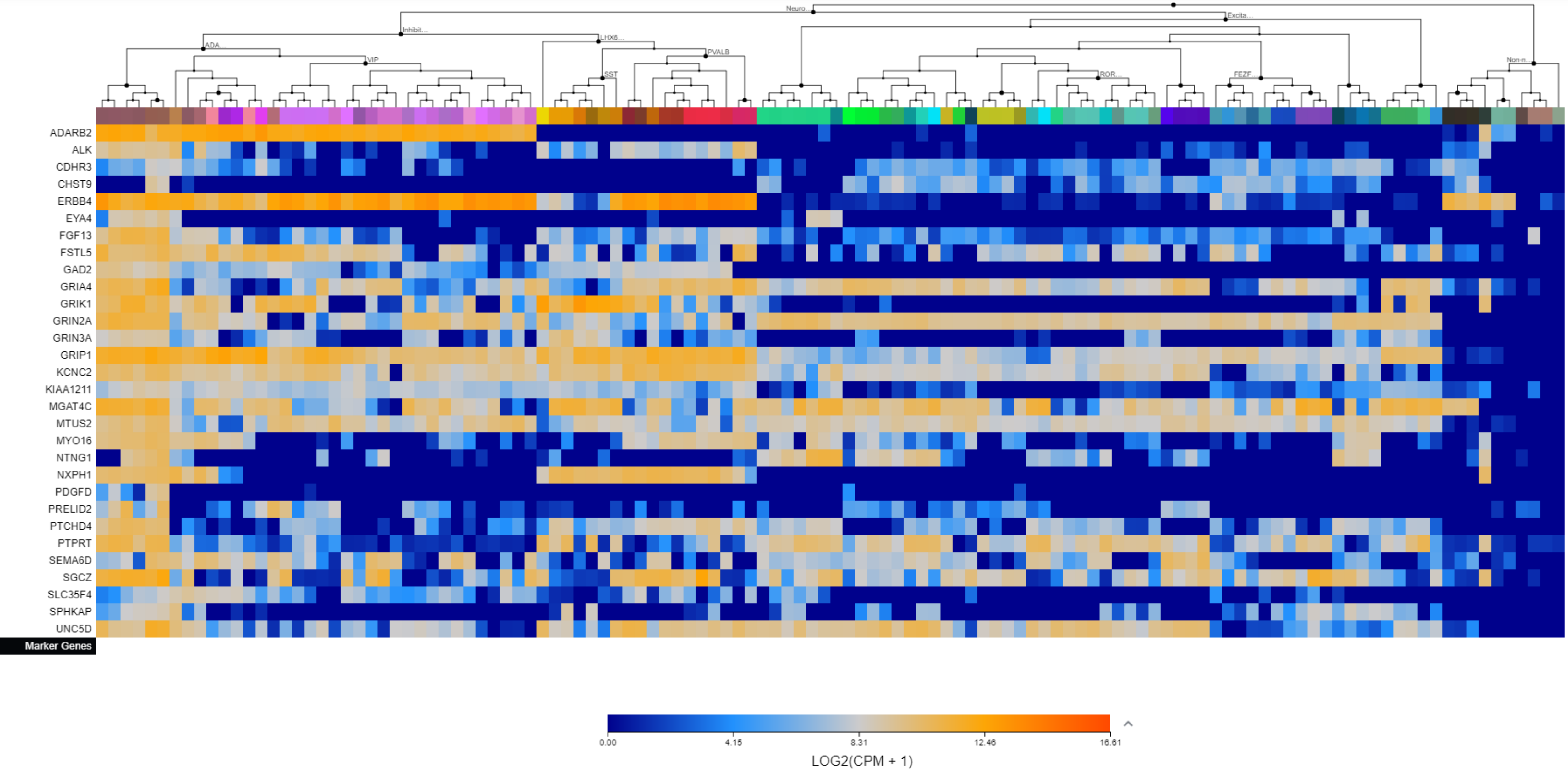

Supplementary File 3. The Allen Brain Map cell type heatmaps by cluster.

Cluster 36

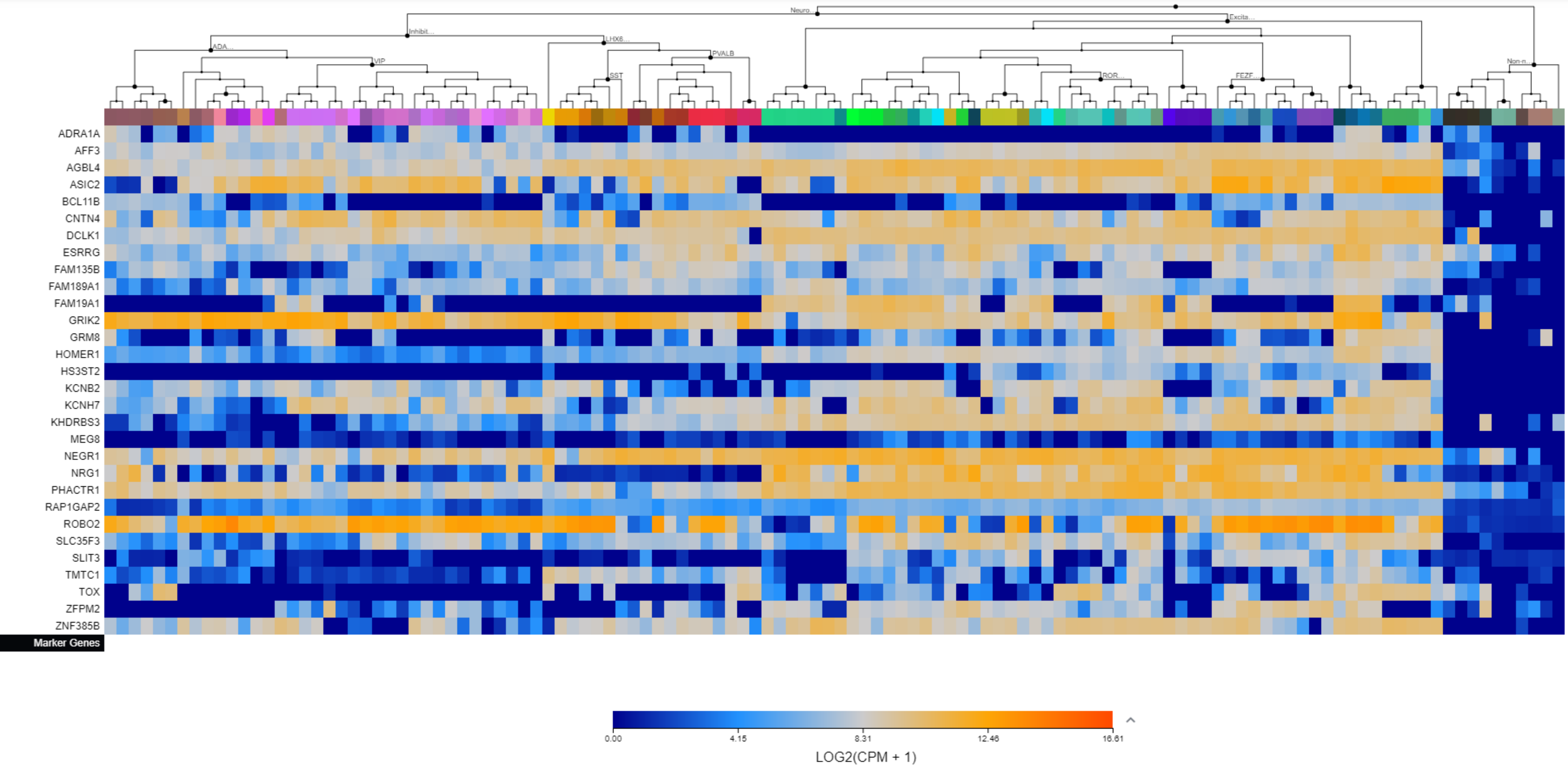

Supplementary File 3. The Allen Brain Map cell type heatmaps by cluster.

Cluster 37

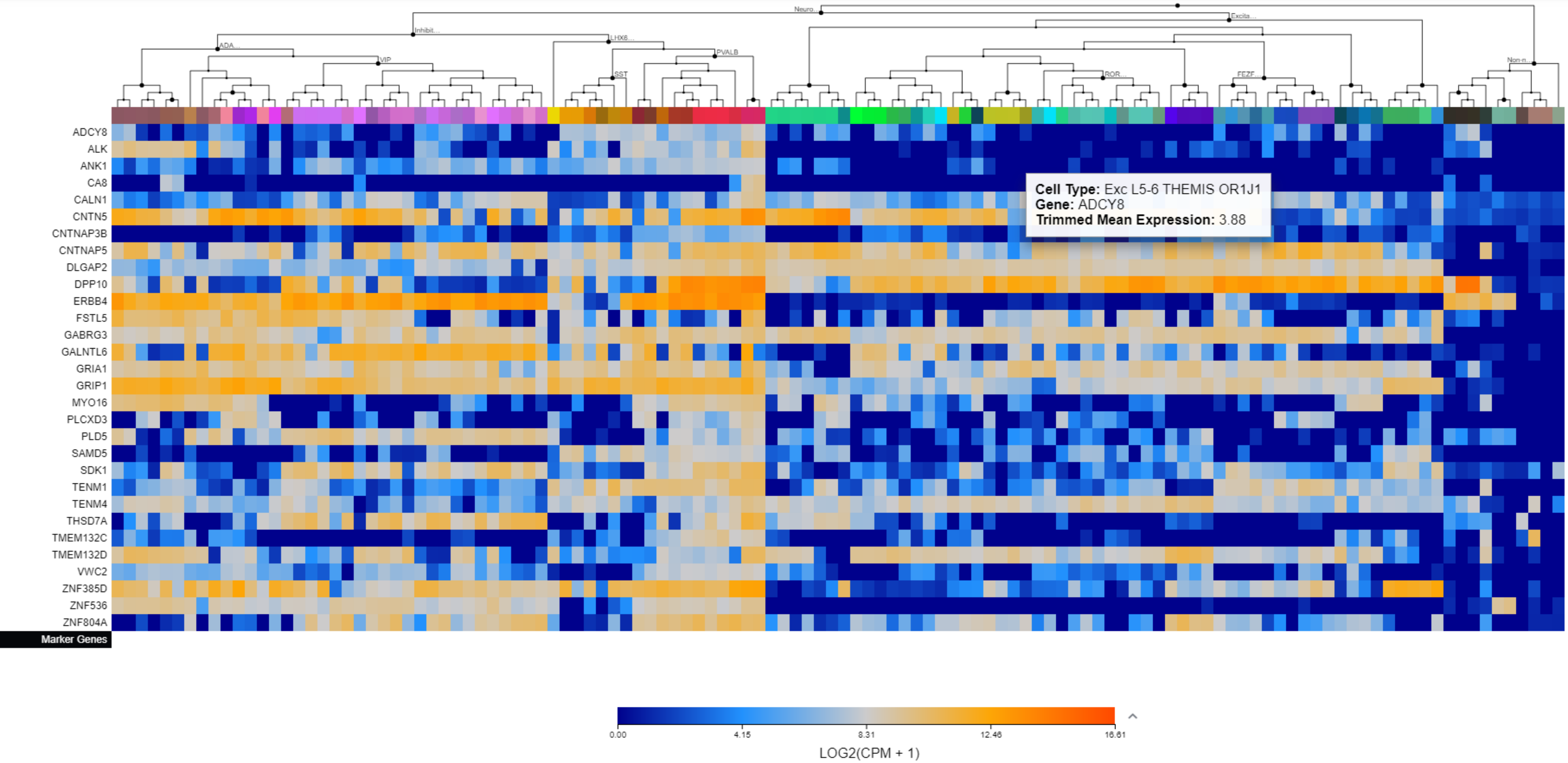

Supplementary File 3. The Allen Brain Map cell type heatmaps by cluster.

Cluster 38

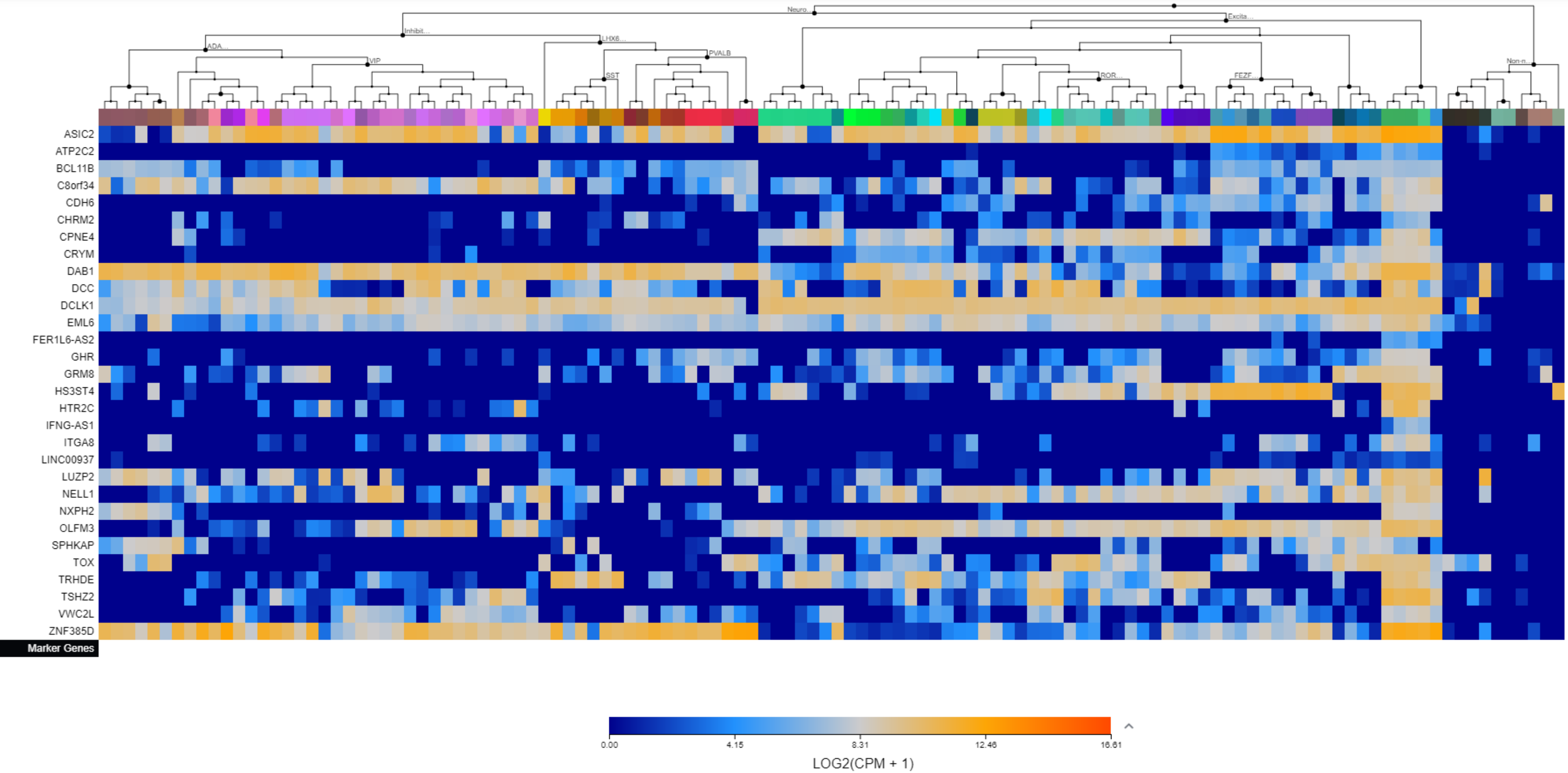

Supplementary File 3. The Allen Brain Map cell type heatmaps by cluster.

Cluster 39

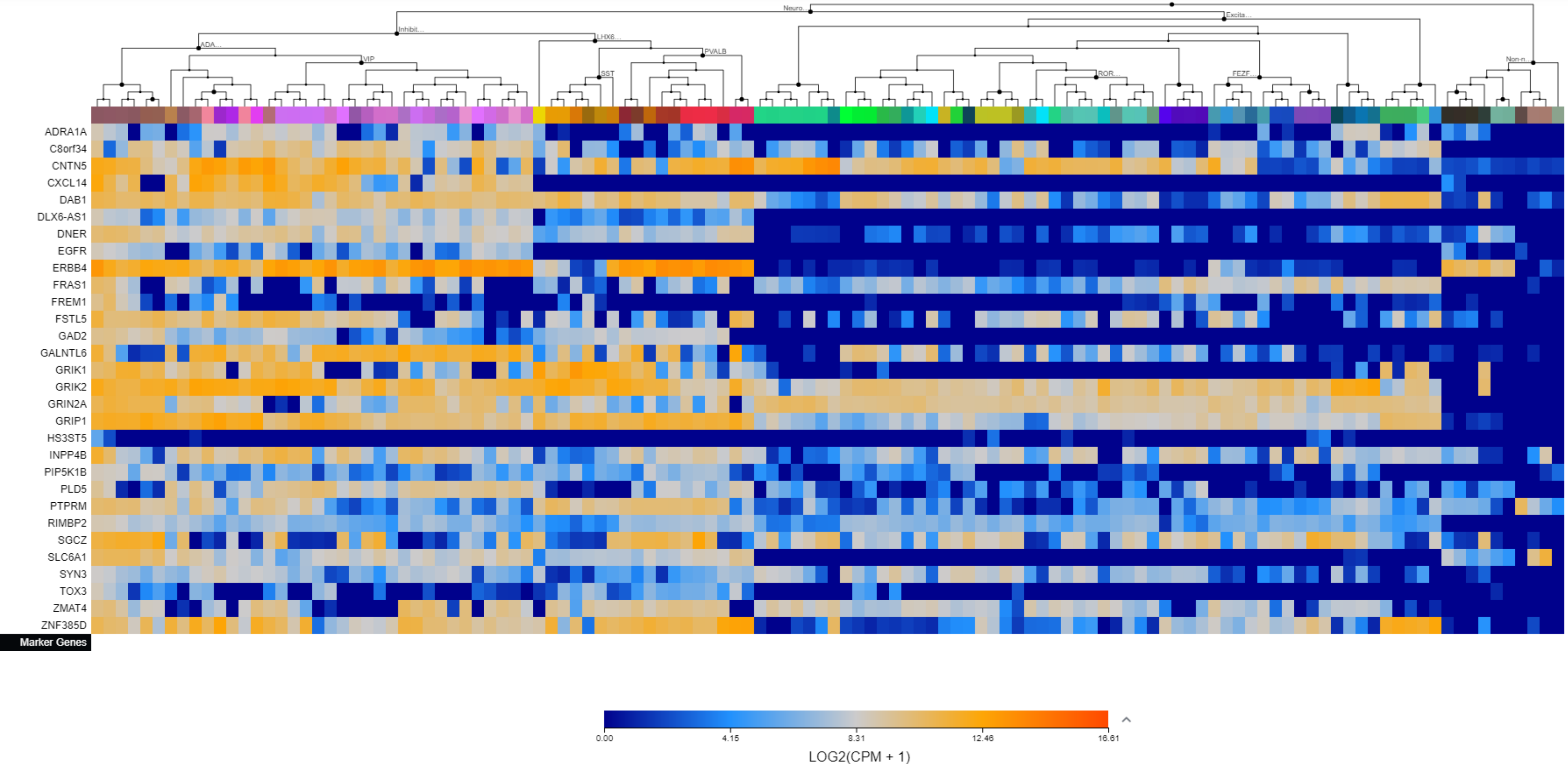

Supplementary File 3. The Allen Brain Map cell type heatmaps by cluster.

Cluster 40

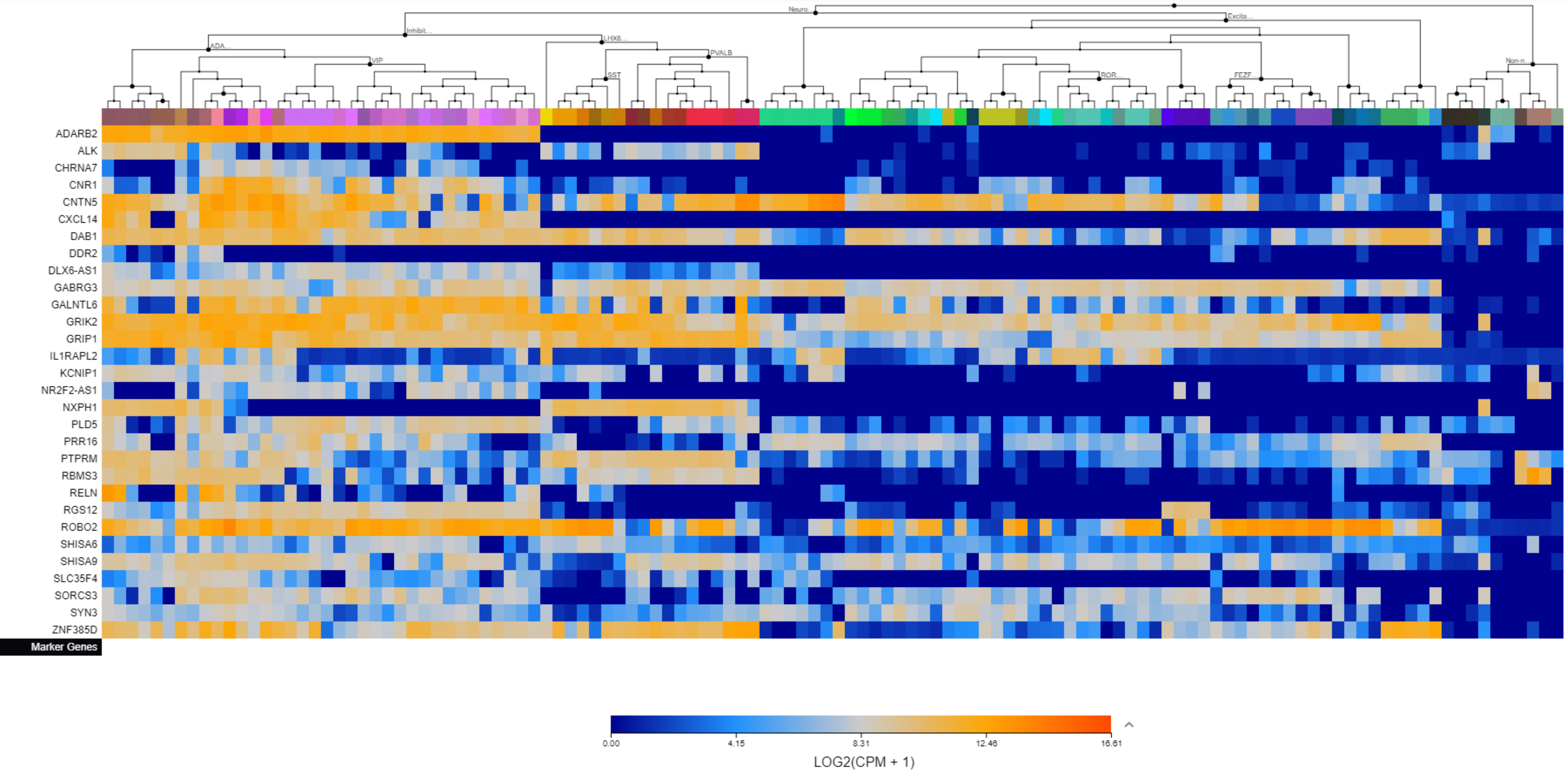

Supplementary File 3. The Allen Brain Map cell type heatmaps by cluster.

Cluster 41

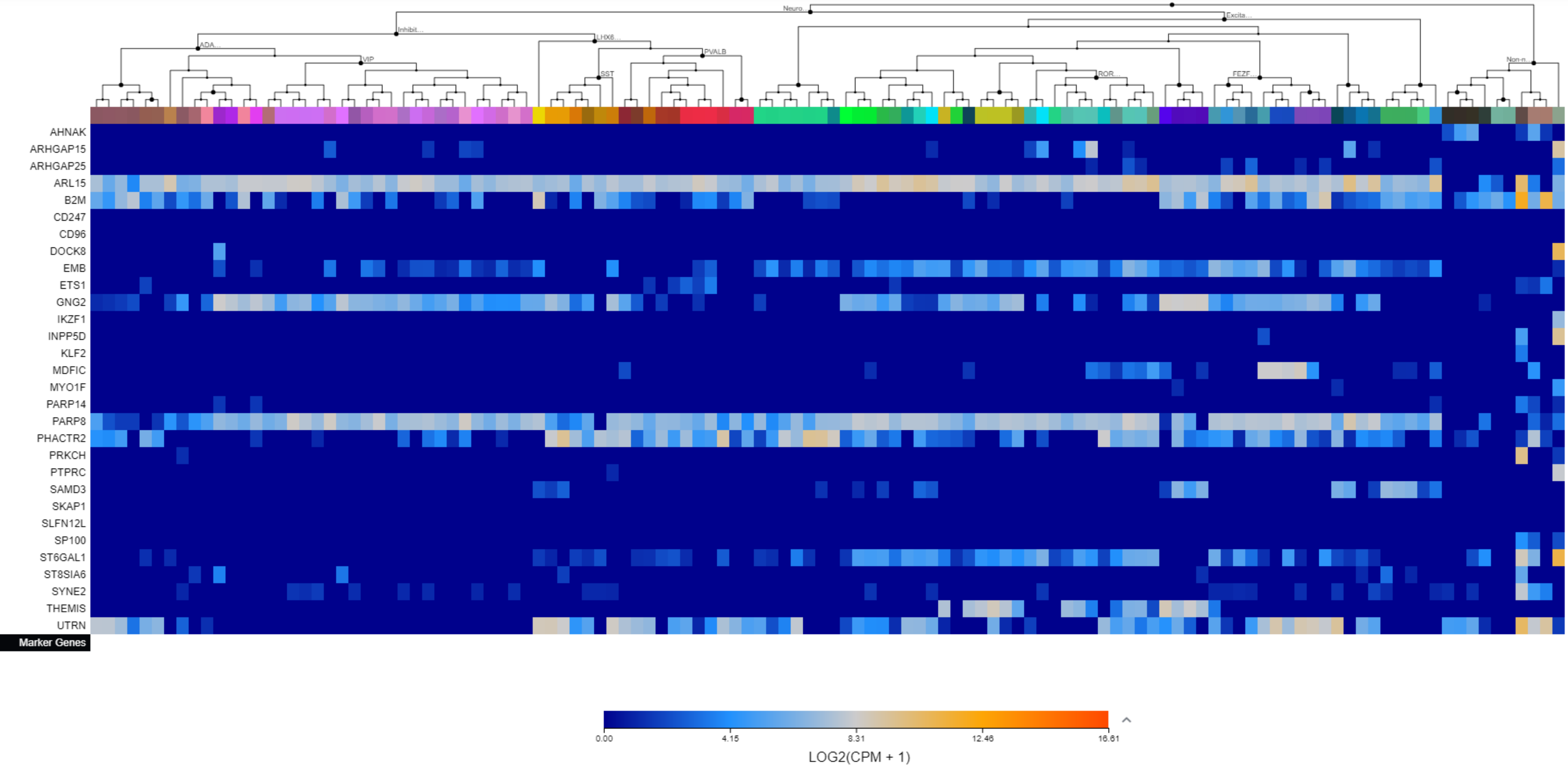
